# Living off the grid: Spatial representations show systematic non-Euclidean distortions regardless of their age and how measured

**DOI:** 10.64898/2026.02.10.705053

**Authors:** Derek J. Huffman, Arne D. Ekstrom, Nikhil Jaha

**Author notes:** Correspondence for this paper should be addressed to Derek J. Huffman, 5568 Mayflower Hill Drive, Waterville, ME 04901.

## Abstract

Spatial memory is invaluable for most mobile organisms, yet nature of the underlying representations that we employ for spatial memory has been fiercely contested. On the one hand, the presence of place cells in the hippocampus and grid cells in the medial entorhinal cortex appear to support the argument spatial representations may follow Euclidean axioms, termed “the cognitive map hypothesis.” On the other hand, decades of behavioral research in humans reveals that spatial memory often shows characteristic distortions, leading to the alternative, cognitive graph hypothesis, to account for this aspect of spatial memory. Importantly, the majority of laboratory studies tend to occur within novel environments in which participants often have only limited exposure and no personal relevance. We were interested in studying large-scale memory across multiple time scales: from a virtual environment (e.g., learned over several minutes) to a college campus (e.g., months to a few years) to a hometown environment (e.g., many years). Across several tasks, we found that participants exhibited systematic distortions in their memory for all of these environments. Likewise, we found significant correlations between performance on several spatial memory tasks (both between participants and within-participant analyses of patterns of errors), thus suggesting that these tasks tap into partially overlapping cognitive representations and supporting their construct validity. Altogether, our findings provide clear evidence for cognitive graph hypothesis and support the construct validity of several spatial memory tasks within large-scale, real-world environments that are learned over the course of several months to years.

**Public Significance Statement:** Spatial memory is key for our ability to live independent lives (e.g., patients with Alzheimer’s disease lose independence, partially due to disorientation in familiar environments). Typical laboratory-based measures of spatial memory use novel environments that may differ in complexity vs. real-world environments (e.g., size, layout, number of landmarks, duration of exploration, personal relevance). We leveraged breakthroughs in technology to study spatial memory across several tasks and temporal scales, from a novel environment navigated over the course of several minutes to a university campus (e.g., months to years) to hometowns (e.g., years to decades). We observed consistent evidence for systematic distortions in spatial memory, which supports the hypothesis that spatial memory is supported by a cognitive graph, thus posing an important challenge to the extremely influential Euclidean, “cognitive map” hypothesis that was awarded a Nobel Prize in 2014.

---

In our daily lives, we commonly navigate over a variety of scales of spaces, from small-scale vista spaces (e.g., rooms in which all parts of the room are perceivable without ambulation) to large-scale environmental spaces (e.g., walking in a wooded forest or navigating cities with barriers occluding direct visibility; Montello, 1993). Prominent theories have suggested that these different scales of space might recruit different cognitive and neural mechanisms (e.g., Brunec, Bellana, et al., 2018; Ekstrom & Isham, 2017; Ginosar et al., 2023; Meilinger, 2008; Meilinger et al., 2014; Montello, 1993; Peer et al., 2019; Wolbers & Hegarty, 2010; Wolbers & Wiener, 2014). The fields of spatial cognition and navigational neuroscience have taught us much about the potential mechanisms of spatial memory (e.g., Ekstrom et al., 2018; Epstein et al., 2017; Moser & Moser, 2008; O’Keefe & Nadel, 1978; Taube, 2007; Waller & Nadel, 2013); however, the dominant paradigm consists of either asking participants (e.g., humans, rats) to learn small-scale spaces (with real-world and virtual environments), to learn spatial environments over a short period of time (e.g., minutes to hours of exploration), or to learn virtual environments (e.g., via a computer’s desktop screen or immersive virtual reality [VR]). Moreover, the environments that researchers employ in traditional experiments typically have little to no personal relevance to the participant. Therefore, we were interested in determining the nature of spatial memory for large-scale, real-world spaces that are learned over long periods of time and for environments that are personally relevant for the participant. We were also interested in how these real-world large-scale spaces would compare with more traditional experimental frameworks, like novel virtual environments.

The nature of the spatial representations that we form when we navigate remains unclear and is a topic of current debate in the literature (Bellmund et al., 2018; Chrastil & Warren, 2014; Du et al., 2023; Ekstrom et al., 2017; Montello, 1993; Moser & Moser, 2008; O’Keefe & Nadel, 1978; Warren, 2019; Warren et al., 2017). According to one set of extremely influential theoretical proposals that fall under the broad idea of the “cognitive map” (e.g., including the 2014 Nobel Prize for the discovery of place cells and grid cells: Hafting et al., 2005; O’Keefe & Dostrovsky, 1971), the representations that we form should generally resemble those of the physical space that we navigate, and therefore obey “Euclidean” axioms (Bellmund et al., 2018; Gallistel, 1990; Moser & Moser, 2008; O’Keefe & Nadel, 1978). As such, the maps that we draw, particularly from cities that we know well, although subject to some degree of error (e.g., global stretching and shrinking of landmarks within the city), should show little, if any, systematic distortions (e.g., finding systematic warping based on environmental features, such as alignment heuristics, would challenge the Euclidean hypothesis). In other words, across landmarks and participants, spatial memory errors should be normally distributed around a mean of the correct locations (e.g., Bonin & Huffman, 2026). In contrast, the labeled graph hypothesis and other “Non-Euclidean” models, suggest that spatial memory for landmarks may be stretched or compressed, thus compromising the global structure of the environment (Chrastil & Warren, 2014; Du et al., 2023; Ericson & Warren, 2020; Warren, 2019; Warren et al., 2017). Such models would predict systematic distortions in spatial memory and would be more akin to a subway map (i.e., topologically accurate) rather than a globally consistent cartographic map (i.e., which would be metrically accurate in a globally consistent manner).

Several studies report distortions in spatial memory that involve non-Euclidean spatial reconstructions when participants remember familiar spatial layouts, for example, violating the property of triangle inequality (Moar & Bower, 1983) or symmetry (Mcnamara & Diwadkar, 1997). Other studies have developed non-Euclidean virtual environments in which participants navigate on environments that loop back on themselves in ways that violate physical reality (Du et al., 2023; Ericson & Warren, 2020; Muryy & Glennerster, 2018, 2021), including wormholes (Warren, 2019; Warren et al., 2017). For example, when participants learned environments in highly immersive VR that contained wormholes that teleported them to a distinct location, the wormholes induced significant non-Euclidean distortions into their navigation and memory judgments despite the fact that participants were unaware of the wormholes’ presence (Warren et al., 2017). Another series of studies showed that when participants walked hallways that looped back on themselves, they distorted the lengths of the hallways to accommodate either missing or impossible intersections (Du et al., 2023). One study, however, reported that participants selected a Euclidean coordinate system when navigating compared to other options, like a spherical or hyperbolic coordinate system (Widdowson & Wang, 2022).

Importantly, though, all of these studies were conducted in virtual reality and involved mismatches between what the participant saw and how they physically ambulated the environment (i.e., a mismatch between visual information and body-based cues) and none of these studies involved well-learned, personally relevant environments with significant exposure (e.g., months to years of exploration). Therefore, it is unclear if non-Euclidean distortions also occur in well-learned, real-world environments.

Why might previous studies have reported non-Euclidean spatial distortions? Navigation behavior is often imperfect, particularly when an environment is not well learned, and thus has been argued to be supported by heuristics or non-Euclidean representations of space in many instances as opposed to a cognitive map (Bennett, 1996; Chrastil & Warren, 2014; McNamara, 1986, 1991; Stevens & Coupe, 1978; Tversky, 1981, 1992, 1993; Warren, 2019). Some participants perform notoriously poorly on spatial memory tasks, including pointing tasks, with some otherwise healthy participants performing at or near chance levels (e.g., Chrastil & Warren, 2013; Foo et al., 2005; Ishikawa & Montello, 2006; Meilinger et al., 2014; Waller & Greenauer, 2007; Weisberg et al., 2014). It is possible that some of the non-Euclidean distortions reported in previous studies might have arisen due to poorly learned, unstable, or changing spatial representations rather than non-Euclidean organization, per se (e.g., perhaps participants did not have ample exposure to the environment to form representations that are supported by Euclidean axioms). Therefore, testing participants in highly familiar environments is important to allow us to determine whether such non-Euclidean distortions are a more general property of human memory or whether a Euclidean model would be favored in these situations. It is also possible that some of the non-Euclidean distortions reported in previous studies manifested due to the specific manner in which participants were tested, with most studies involving either pointing or walking to unseen targets as some manipulation was performed in virtual reality (i.e., many previous studies did not compare if there were consistent distortions across different spatial memory measures). Thus, we argue that it is important to test participants with several different dependent measures, including map drawing and pointing to determine whether such tasks elicit similar error patterns.

Map drawing is often taken as the “gold-standard” for allocentric forms of spatial representations, although these are often the most difficult for participants to acquire. Indeed, when learning an environment from a map, participants often assume that “up” on the map corresponds to the forward direction when placed within the environment, and this alignment heuristic causes performance to suffer dramatically when participant’s orientation within the environment is misaligned with that of the map (Gagnon et al., 2014; McNamara et al., 2003; Mou & McNamara, 2002; Richardson et al., 1999; Shelton & McNamara, 1997, 2001). Another manifestation of such non-Euclidean biases in how people remember spatial locations is termed an alignment effect, which are commonly observed in pointing judgments in the judgments of relative direction (JRD) task (“Imagine you are standing at A, facing B. Please point to C”; Ekstrom et al., 2017; Huffman & Ekstrom, 2019; Rieser, 1989; Shelton & McNamara, 2001, 2004; Waller & Hodgson, 2006). These manifest most often in tests of pointing accuracy to hidden locations such that participants tend to remember locations “aligned” with the principle axes of the environment (such as defined by a boundary) better than those misaligned (see Figure 1B; Frankenstein et al., 2012; Mou et al., 2006, 2007; Shelton & McNamara, 2001, 2004). Whether alignment effects manifest with map drawing is less clear. Alignment effects are a form of non-Euclidean distortion because they involve remembering aspects of the environment in a way that is systematically distorted: landmarks are remembered better that align with the boundaries or cardinal direction of the environment than those that do not. Such alignment effects, though, have not been connected to maps that are drawn and how consistent these across familiar, recently learned, and novel environments is unclear (Starrett et al., 2019), particularly in large-scale environments that lack clear “boundaries.” One study that looked at maps drawn of towns of some participants suggested non-Euclidean distortions, although these cities were not experimentally manipulated in terms of levels of familiarity, how recently they were experienced, overall accuracy, nor how this related to other spatial memory measures, such as pointing accuracy and distance estimation (Nakaya, 1997). Thus, it is important to resolve whether well-learned cities, compared to novel cities, show distortions in how they are structured and how these might vary depending on the dependent measure.

**Figure 1:**
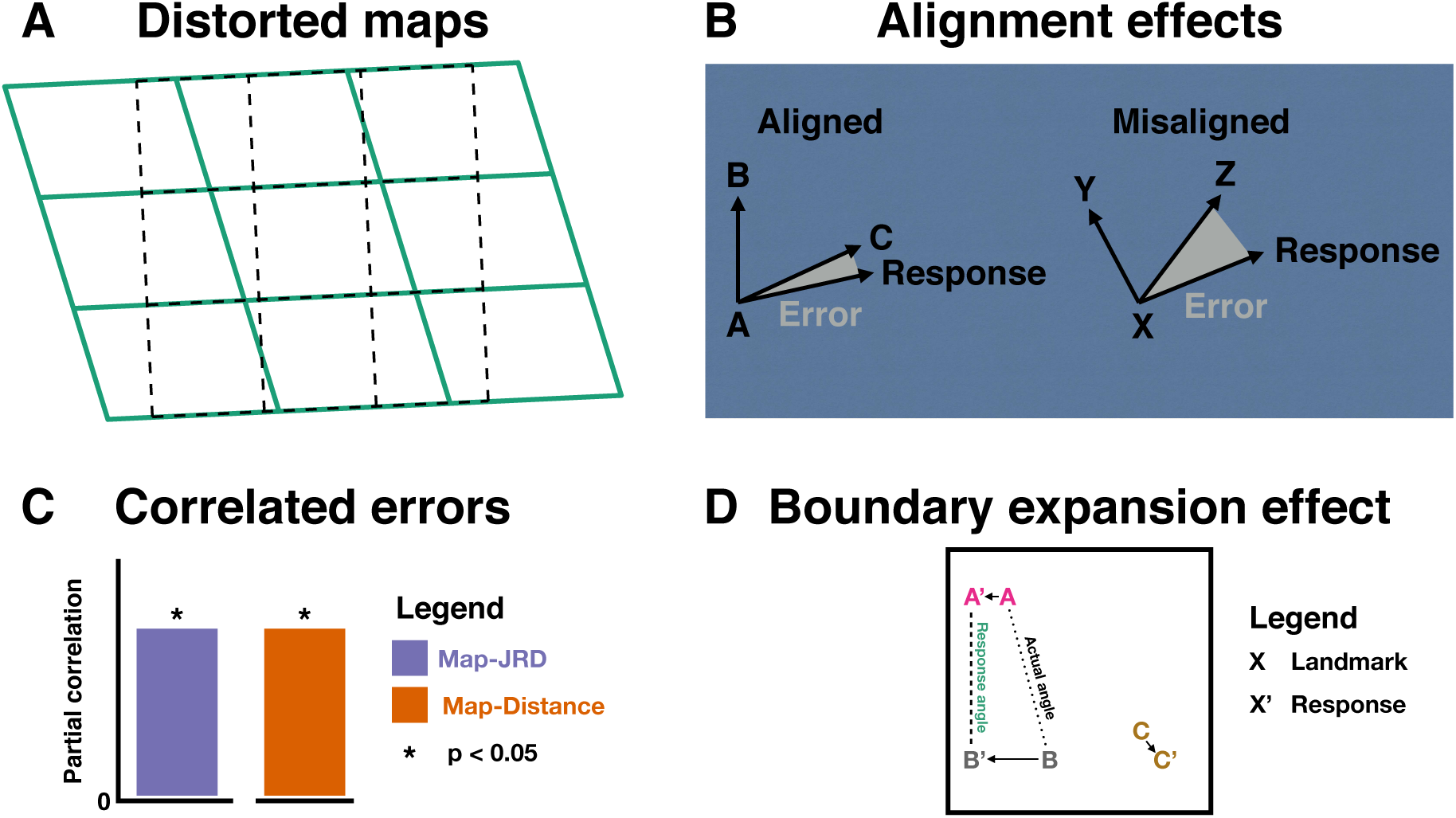
Overview of our predictions and main findings, which all support the hypothesis that the cognitive representations supporting human spatial memory are guided by systematic distortions. A) We found that Map Drawing Performance for the real-world environments was better explained by an affine transformation (green solid lines) than a Euclidean transformation (black dashed lines; note the significant shearing and distortions here, which depict the affine transformation for the participant with the median ΔAIC in favor of the affine model for the Hometown Experiment). B) We found evidence for boundary alignment effects in pointing performance on the JRD task (i.e., participants pointed better for imagined headings that were aligned with the salient axes of the environment than misaligned) for both real-world environments and the virtual environment. C) We found that the pattern of errors was correlated between the Map Drawing Task and both the JRD task and the Distance Estimation task for both real-world environments and the virtual environment, thus suggesting that participants recruit similar underlying representations to solve these tasks. D) We found evidence for a boundary expansion effect in which participants placed the landmarks closer to the boundaries than they actually were in their map drawing data for the virtual environment, which can also help explain the observed alignment effects.

Here, we test four key predictions of the cognitive graph hypothesis (Figure 1). To test participants on familiar environments, we used Google Maps to assess the accuracy and nature of distortions in their map drawing and pointing responses. We also included a novel virtual environment that we built in the Unity Game Engine that had a similar structure to real-world environments. To test for distortions, we looked at both how landmarks within the environment were either aligned or misaligned with the global axes and whether a Euclidean or affine (allowing more distortions, such as an alignment distortion) best described their drawn maps. Our first main prediction of the cognitive graph hypothesis is that participants’ map drawing performance for well learned, real-world environments will be significantly better fit by an affine transformation than a so-called Euclidean transformation, which would suggest that participants maps are significantly distorted, which is consistent with the use of alignment and rotation heuristics (see Figure 1A). Our second main prediction from the cognitive graph hypothesis is that participants’ JRD responses will exhibit an alignment effect for all of the environments, which would provide additional support for an alignment heuristic (see Figure 1B). Our third main prediction is that we will observe correlations between performance and patterns of errors across the JRD Task, the Map Drawing Task, and the Distance Task across all environments, which would suggest that these tasks tap into partially overlapping representations, and further demonstrate the ubiquity of our findings across tasks (see Figure 1C). Our fourth prediction from the cognitive graph hypothesis is that participants’ maps for the virtual environment will exhibit clear distortions. To foreshadow the specific distortions in these maps, we found a “boundary expansion effect” and an alignment effect of landmarks along the boundaries of the environment (see Figure 1D). Altogether, our approach is important because will allow us to discover whether the non-Euclidean distortions observed previously for novel virtual environments replicate with novel environments that do not involve visual-motor mismatches performed in VR and further extend to even well-learned, real-world environments that participants have extensively experienced. Moreover, tested whether we could provide evidence for systematic distortions in both map drawing and pointing, which would suggest that our results are insensitive to testing method. Alternatively, the highly influential Euclidean, cognitive map hypothesis would predict that we would not observe evidence of systematic distortions across these measures. To foreshadow our conclusion, our results provide strong support for the cognitive graph hypothesis across tasks and environments.

## 2 General Methods

## 2.1 Participants

Participants in both experiments were University of California, Davis students and they received course credit for their participation. All participants consented to participation in accordance with the Institutional Review Board of University of California, Davis. All data collection took place in person over the course of two sessions. We used a large testing bay for the sessions, allowing us to test up to 15 participants at a time, thus allowing for the relatively large samples in both of our experiments.

## 2.2 Tasks

In both experiments, we tested spatial memory for a familiar real-world, large-scale environment, which differed between our two experiments (see below). Briefly, in Experiment 1, we tested participants’ knowledge of the University of California, Davis campus, while in Experiment 2 we tested participants’ knowledge for their hometown. Importantly, the UC Davis campus is designed such that cars cannot drive through the main part of campus; thus, the representations that participants form likely include information from body-based cues (especially those from biking [a very common mode of transportation in Davis] and walking around campus). We also assessed participants’ ability to form new spatial memories for virtual environments using a desktop-based virtual navigation task.

### 2.2.1 Virtual environment learning

Participants learned about a virtual environment that contained 10 landmarks. The virtual environment and the navigation task were the same as in our previous work. Specifically, as we describe in previous work (Huffman & Ekstrom, 2019), we programmed our experiment and our environment in the Unity game engine (https://unity3d.com). Our virtual environment was a 200 × 200 virtual meters (vm) open, square arena (i.e., a large-scale vista space). The boundaries of the environment were brick walls (6 vm tall). The environment contained 10 landmarks (i.e., stores), and we selected the coordinates of the stores such that the distribution of the imagined headings relative to the main axes of the environment was approximately uniform (calculated with custom-written R code). For each participant, the 10 stores were randomly selected from a list of 23 stores—in alphabetical order: 1st Bank, Bakery, Barber, Bike Shop, Book Store, Butcher Shop, Camera Store, Cell Phones Store, Chinese Food Restaurant, Clothing Store, Coffee Shop, Craft Shop, Dentist, Fast Food Restaurant, Florist, Grocery Store, Gym, Ice Cream Shop, Music Store, Pet Store, Pharmacy, Pizzeria, and Toy Store—and each store was randomly assigned to one of the 10 landmark locations. The random assignment of stores controls for semantic effects of the stores across participants. Each store was 10.9 vm long × 15 vm wide × 6 vm tall and differed in the colors of the walls and awnings as well as unique objects in the windows. The eye height of the avatar was 1.56 vm above the ground.

Before exploring the virtual environment, participants viewed each of the 10 landmarks for 3 seconds. We instructed participants to read the name of the store and to try to memorize its name and visual features. Next, we asked participants to navigate to each of the 10 landmark stores in a random order. Participants indicated that they found each store by walking into the walls of the store, which would then tell the participant the name of the next store that they need to find. After participants found each of the 10 stores, they performed the judgments of relative direction (JRD) task. They performed 6 interspersed blocks of navigation and the JRD task before the other spatial memory tasks.

### 2.2.2 The judgments of relative direction task

During the Judgments of Relative Direction (JRD) Task, we asked participants to point in the location of landmarks in the environment. Specifically, each trial began with the following prompt: “Imagine you’re standing at X.” We instructed participants to respond, by pressing the space bar, as soon as they could imagine themselves standing at X. Then, the following prompt was added to the screen: “Facing Y.” Again, we instructed participants to respond as soon as they could imagine themselves facing Y. Then, the following prompt was added to the screen: “Please point to Z.” Again, we instructed participants to respond as soon as they could imagine the angular response that they were going to make on the trial. Then, the participants saw a compass and arrow appear on the screen and they used the arrow keys to rotate the arrow to match their memory of the relative direction for that trial. The benefit of the separate responses to the standing, facing, pointing, and the response phase is that it would allow us to look at the relative reaction times of these various aspects of the task.

### 2.2.3 The distance estimation task

During the Distance Estimation Task, we asked participants to imagine the straight-line distance (i.e., the beeline distance or the distance as the crow flies) between landmarks in the environment. Each trial of the distance task began with an instruction for the participant to imagine a given standing location. We instructed the participants to respond, by pressing the space bar, as soon as they could imagine themselves standing in that location. Then, the participant viewed the name of a second location. We instructed participants to respond as soon as they knew their estimate of the distance from the imagined standing location to the location of the second landmark. Then, participants input their responses by typing numbers, which showed up on the screen. When participants were done typing text for that trial, they pressed Return/Enter to submit their response.

### 2.2.4 The map-drawing task

We also asked participants to draw a map of the environment using a model-building task. Specifically, participants clicked and dragged virtual objects around on the screen to create a map (i.e., viewed from overhead). For the virtual environments, the task began by showing an overhead view of the environment, including the bounding walls of the environment, with all of the landmarks placed in a straight line (top to bottom) outside of the environment (the assignment of each landmark to the 10 starting coordinates was randomized for each participant). Then, participants clicked and dragged the stores into the environment. We only allowed participants to submit their responses if they had moved all of the stores within the bounding walls of the environment.

The Map-Drawing Task was very similar for the real-world tasks. However, instead of using virtual buildings, etc., the participants saw a circle with a text label above each circle corresponding to the name of a landmark. Participants then clicked and dragged the circle/text around on the screen to generate their maps. Here, participants had to move every circle/text box to be able to submit their final responses.

### 2.2.5 The recall task

We had participants perform a recall task similar to previous work (Hirtle & Jonides, 1985; McNamara et al., 1989); however, due to space constraints, we do not discuss the data here.

## 2.3 Calculating angles and distances for our real-world tasks

In our real-world experiments, we used Google’s “My Maps” to gather location information for our landmarks. In Experiment 1, we manually selected the center of the campus buildings. In Experiment 2, our participants manually selected the locations (i.e., from their hometowns) at the end of the session. These maps provide longitude and latitude coordinates in WGS84. We loaded these maps into R using the package sf (version 0.8-1). To calculate angles between landmarks, we calculated the initial bearing between these coordinates using the function *bearing* from the R package geosphere (version 1.5-10). To calculate distances between landmarks, we calculated the geodesic using *distGeo* from geosphere. To analyze map data (e.g., our bidimensional regression analysis between the actual locations and participants’ responses), we projected the coordinates to the Universal Transverse Mercator using *st_transform* from sf.

## 2.4 Analyzing performance on the Distance Estimation Task

To analyze performance on the Distance Estimation Task, we calculated the Spearman rank correlation coefficient between participants’ distance estimates and the answers (calculated as the Euclidean distance for the virtual environment and the geodesic for the real-world experiments; see 2.3). Note, we used the Spearman rank correlation coefficient (instead of the Pearson correlation coefficient) because previous research has suggested that the relationship between subjective and objective distances exhibit a monotonic but non-linear relationship (e.g., a power function; Thorndyke, 1981). Therefore, here we are assuming that stronger correlations between the participant’s responses and the answers indicate better performance on the distance estimation task, and our approach obviates the need for having participants make judgments in explicit units (i.e., we are measuring relative distances).

## 2.5 Investigating the relationship between performance on the JRD Task and the Map Task

To test whether there was a relationship between participants’ pointing performance on the JRD Task and the Map Task, we used a partial correlation approach to assess the similarity of pointing angles between the JRD Task and the Map Task while partialing out the relationship between these angles and the answers. This method effectively calculates the similarity between the patterns of errors on these two tasks (note: in previous work we used a slightly different approach, Huffman & Ekstrom, 2019); however, simulations revealed that the partial correlation method is preferred over methods such as correlating the error patterns for angular data (see Appendix). We used the following equation to calculate partial correlations (Krzanowski, 2000):

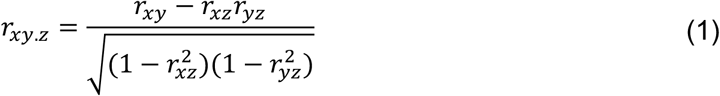

where *x* represents the circular array of angular responses on the JRD Task, *y* represents the circular array of angular responses on the Map Task, *z* represents the circular array of angular answers for each of the questions on the JRD Task, and *r* represents the circular correlation coefficient (calculated via *cor.circular* from the R package circular [version 0.4-93]).

## 2.6 Investigating the relationship between performance on the Distance Estimation Task and the Map Drawing Task

We used a similar within-subjects approach to what is described above for the JRD Task and Map Drawing Task except for the Distance Estimation Task and the Map Drawing Task (i.e., we used the distance estimates from both the Distance Estimation Task and the Map Drawing Task and we partialed out the effect of the actual distance between the points using Equation 1).

## 2.7 Investigating alignment effects

To test whether there were alignment effects (Marchette et al., 2011; McNamara et al., 2003; Mou & McNamara, 2002; Shelton & McNamara, 1997, 2001), we employed a sinusoidal regression. For our experiments that used the spatial environments across participants (i.e., the virtual environment for all of our experiments and the real-world environment for Experiment 1) we first determined the salient axes of the environment. For the virtual environment, this corresponded to the walls of the environment, where we assumed that the error would be maximal at the axes that were offset from the axes of the environment by 45 degrees. For the UC Davis campus, we used OSMnx to calculate the street orientations (Boeing, 2017). We then subjected these values to a sinusoidal regression:

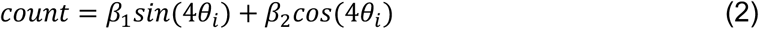

where *count* indicates the number of streets within each given angle bin (denoted by *θ_i_*). We then shifted the angle by 45 degrees because alignment effects would predict that error would be lowest when participants were aligned to the preferred angle and maximal when they were 45 degrees from the preferred angle. We then tested whether the environmentally defined preferred angle accounted for significantly more variance (i.e., indicating an alignment effect) relative to a null model:

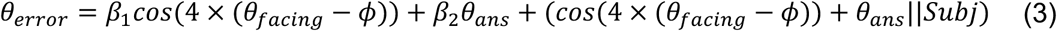

We compared the full model (above) to a null model without the effect of the sinusoidal model:

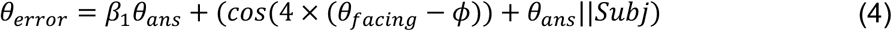

We also employed a version of sinusoidal regression that allowed us to empirically determine the alignment axes. Specifically, for UC Davis campus environment (Experiment 1) we tested a model with a 4-fold pattern:

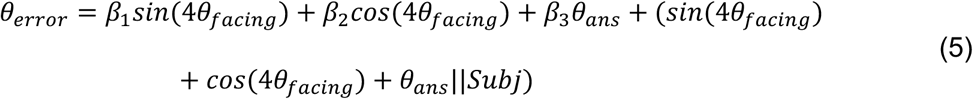

where *θ_error_* is the absolute angular error, *θ_facing_* is the facing angle of the JRD question (in radians), and *Subj* is a factor representing the participant I.D. To account for the possibility that absolute angular error would be modulated by the absolute magnitude of the angular answer (i.e., greater error for pointing directions behind the participant relative to questions with pointing directions in front of the participant), we included a regressor for the absolute magnitude of the answer (here, we converted the angle to absolute angle [i.e., between 0 and 180 degrees] and then z-scored these answers)—this term is represented as *θ_ans_*. Previous studies have implemented similar methods, especially for the study of real-world environments (Marchette et al., 2011). To account for the non-normal nature of the absolute angular error, we fit these models using a generalized linear model framework using the function *glmer* within *lme*4 with a gamma function (with log link). To test significance, we used a *χ*^2^ test to compare our full model to a null model without the fixed effects terms for the facing angle:

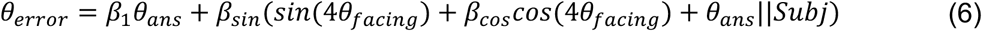

We calculated the preferred angle of the alignment effects using the following equation (e.g., Julian et al., 2018; Nau et al., 2018):

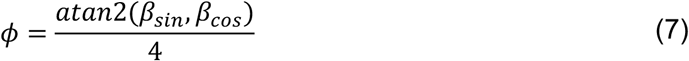

where *β_sin_* and *β_cos_* are the beta coefficients from (5) and *atan*2 was the function from the package base in R. In some cases, we converted these angles from radians to degrees by multiplying *ϕ* by 180*/π*. Note, because we fit our model to the absolute angular error (i.e., greater error), *ϕ* will reflect the offset of maximal angular error and hence the aligned axes will be *π/*4 radians or 45 degrees from this angle (i.e., because this is a measure of error). For example, if the pattern of errors is aligned to the NSEW axes (for the UC Davis campus experiment) or to the boundary walls (the virtual environment in all three experiments), then *ϕ* will be ±*pi/*4 radians or ±45 degrees). Note, for our analysis of real-world locations in Experiment 1 (the UC Davis campus experiment), we calculated angles from the Universal Transverse Mercator projection (see 2.3). For the virtual environment data, we implemented a cosine regression model:

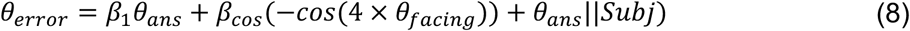

where we included the minus sign because we assumed best performance when the angles were aligned to the main axes of the environment (i.e., the lowest error).

### 2.7.1 Hometown alignment effects analysis

For the Hometown data (Experiment 2), we used a cross-validation approach to account for the fact that each participant had a unique hometown. Specifically, we would expect the orientation of the alignment effects to differ based on the overall alignment of the landmarks, etc. within that city (e.g., some cities are aligned NSEW while others are aligned obliquely and still others are not laid out in a grid at all). Therefore, we ran a four-fold model (similar to the model described above) separately within each participant using the *glm* function of the stats package (version 3.6.3) in R. We implemented an N-fold-cross-validation approach in which we fit the model and tested its fit iteratively within each participant. We fit the model to the training data using a similar approach to our group-level approach described above. We fit a within-subject model with the following equation (using the *glm* function with a gamma function with a log link function):

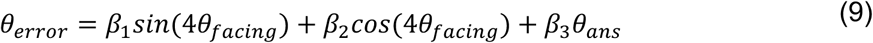

The validation test determine whether a similar alignment effect was found in the left-out data. Our approach is similar to that taken for the analysis of directional coding in fMRI data, and we used the following equation to estimate the fit in our left-out testing data (Julian et al., 2018):

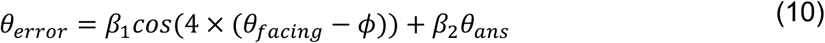

We calculated *ϕ* using Equation 7 and the beta coefficients from the training data using Equation 9. We used a 4-fold cross validation procedure and averaged the resultant beta coefficients for the *cos* term (i.e., *β*_1_) within each participant. To determine whether the 4-fold alignment effect was significant, we tested whether the average beta coefficients were greater than zero at the group level (for a similar approach to fMRI data see: Julian et al., 2018).

## 3 Experiment 1, Real-World Task: UC Davis Campus

### 3.1 Methods

To determine which campus buildings to include in the spatial memory tasks, each participant first performed a landmark confidence task in which they rated their confidence on a scale of 1 to 6 for all 152 campus buildings at University of California, Davis. We then selected the 10 buildings that received the highest confidence ratings. Thus, each participant had a unique set of buildings that were most familiar for them.

### 3.2 Results

#### 3.2.1 Campus task: Evidence that participants’ memories for their local university campus are better explained by distorted representations than by Euclidean maps

We investigated whether participants’ maps of their local university campus were better explained by a representation involving systematic distortions (here, an affine transformation; i.e., translations, rotations, non-uniform scaling, and shearing) than a Euclidean representation (here, a Euclidean transformation; i.e., translations, rotations, and uniform scaling) using bidimensional regression of the actual map locations (i.e., the participants’ responses were the dependent variables and the actual map locations were the independent variables; Carbon, 2013; Friedman & Kohler, 2003; Nakaya, 1997; Tobler, 1994). If participants formed metric, Euclidean representations of their university’s campus, then the Euclidean model should fit the data better than the affine model. Conversely, if participants relied on heuristics (e.g., alignment and rotation heuristics; Tversky, 1981) and formed distorted representations of their college campus, then the affine model should fit the data better than the Euclidean model (cf. Nakaya, 1997). Specifically, an affine transformation allowed for non-uniform scaling and shearing that would be consistent with systematic distortions (e.g., possibly allowing for a rotation or alignment heuristic; Nakaya, 1997). To test these competing hypotheses, we fit each participant’s map data using a Euclidean and an affine bidimensional regression model and we calculated the difference in the Aikake information criterion (ΔAIC) between the model fits (using the R package BiDimRegression [version 2.0.0]; Carbon, 2013). The AIC approach attempts to find the most parsimonious model by penalizing models with more parameters (for application to bidimensional regression see: Nakaya, 1997). Lower AIC values indicate a better fit and a significance threshold is typically set at |ΔAIC| *>* 2 (Nakaya, 1997). We calculated the ΔAIC between the affine model minus the Euclidean model, therefore negative values indicate that the affine model is preferred while positive values indicate that the Euclidean model is preferred.

The ΔAIC was negative for the majority of our participants (the ΔAIC was less than 0 for 68.2% and less than −2 for 50.5% of our participants; see Figure 2). In contrast, the Euclidean model provided a better fit (i.e., ΔAIC *>* 2) in 19.6% of participants and the models were inconclusive (i.e., −2 *<* ΔAIC *<* 2) in 29.5% of participants. Two-tailed Wilcoxon signed rank tests revealed that the median ΔAIC, −2.12, was significantly less than 0 (*p* = 4.66×10^−7^). These results suggest that the structure of participants’ knowledge of their university’s campus is better explained by distorted, heuristic representations than by metric, Euclidean maps (Nakaya, 1997), with some evidence for inter-subject variability (Ishikawa & Montello, 2006; Weisberg et al., 2014).

**Figure 2:**
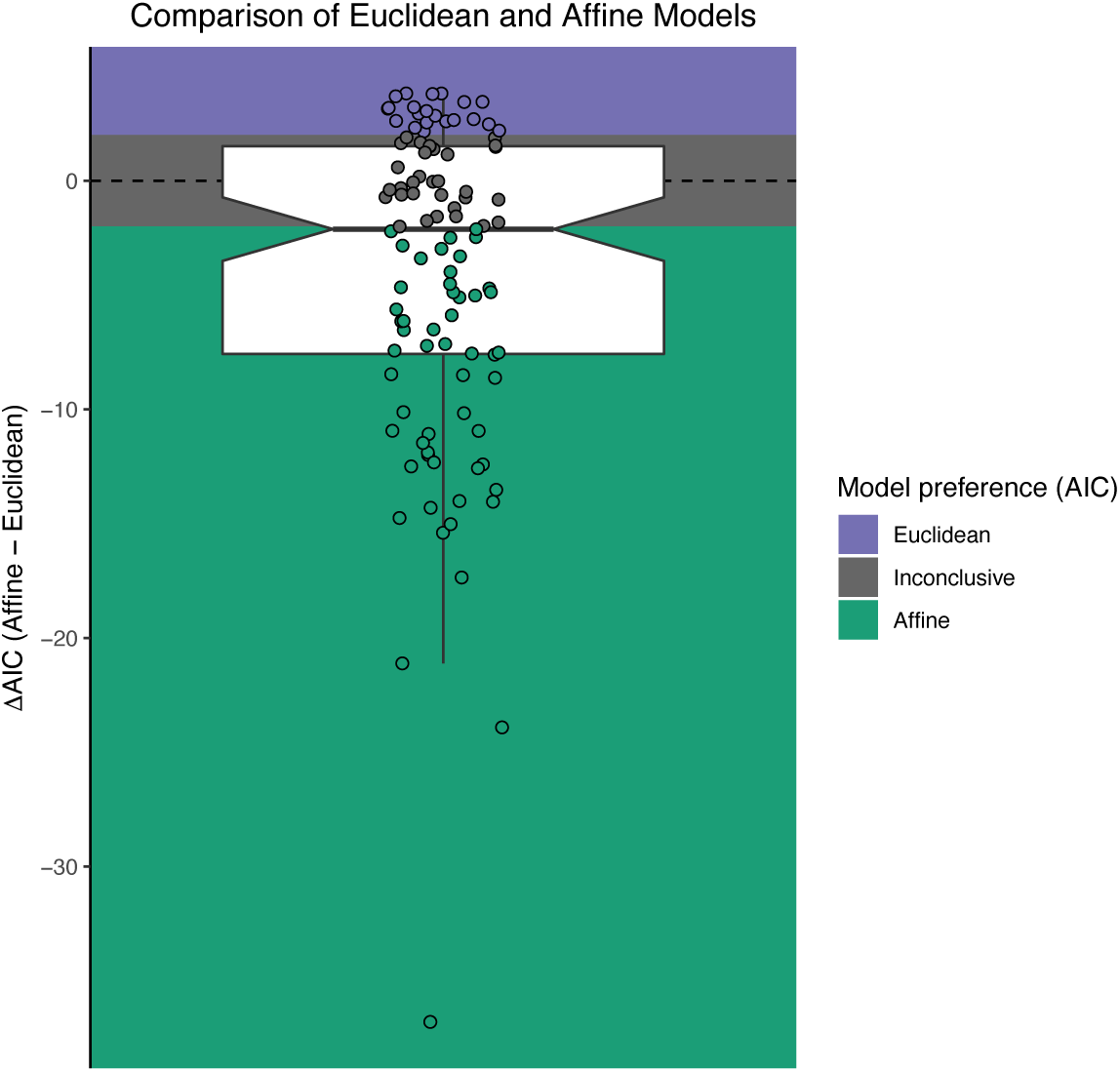
Evidence that participants’ memories for their university campus are systematically distorted. We fit each participant’s map drawing data with a Euclidean and affine transformation. This figure depicts the ΔAIC between the two model fits, where negative values are in favor of the affine model and positive values favor the Euclidean model. The ΔAIC for the majority of participants was less than −2, which is typically taken to be significant evidence in favor of the model. A Wilcoxon signed rank test revealed that the median ΔAIC, −2.12, was significantly less than 0 (*p* = 4.66 × 10^−7^). Each dot indicates an individual participant and the boxplot depicts the middle 25th to 75th percentile of the data. The colors indicate model preference, where the Euclidean model is preferred if ΔAIC *>* 2, the affine model is preferred if ΔAIC *<* −2, and the model comparison is inconclusive if −2 *<* ΔAIC *<* 2.

#### 3.2.2 Campus task: Evidence for Alignment Effects in Spatial Memory

We next tested whether there were alignment effects within the JRD task, which would allow additional testing of non-Euclidean distortions. We compared a null model vs. a sinusoidal regression model with the axes aligned to the main axes of the environment (as calculated by fitting a regression model to the street network map from OSMnx). Here, we found that the street-network alignment model accounted for significantly more variance than the null model (*χ*^2^(1, *N* = 107) = 8.60, *p* = .0034, AIC_null_ = 62,787, AIC_full_ = 62,780, ΔAIC in favor of the full model = 7; see Figure 3). Additionally, when we fit a separate model in which we estimated the best fitting angle of the alignment effects, we found that this model also accounted for significantly more variance than the null model (*χ*^2^(1, *N* = 107) = 8.32, *p* = .016, AIC_null_ = 62,783, AIC_full_ = 62,779, ΔAIC in favor of the full model = 4; see Figure 3). Moreover, the best-fitting empirical angle was remarkably consistent with the salient grid-axes of the environment (within 1.5 degrees, as determined by OSMnx; see the dashed line in the bottom right panel of Figure 3). Altogether, these results demonstrated significant alignment effects thus replicating and extending previous work from small-scale environments (Mou & McNamara, 2002; Shelton & McNamara, 1997, 2001), large-scale and newly learned environments (McNamara et al., 2003), and one other published paper on participants’ memory for their college campus (Marchette et al., 2011).

**Figure 3:**
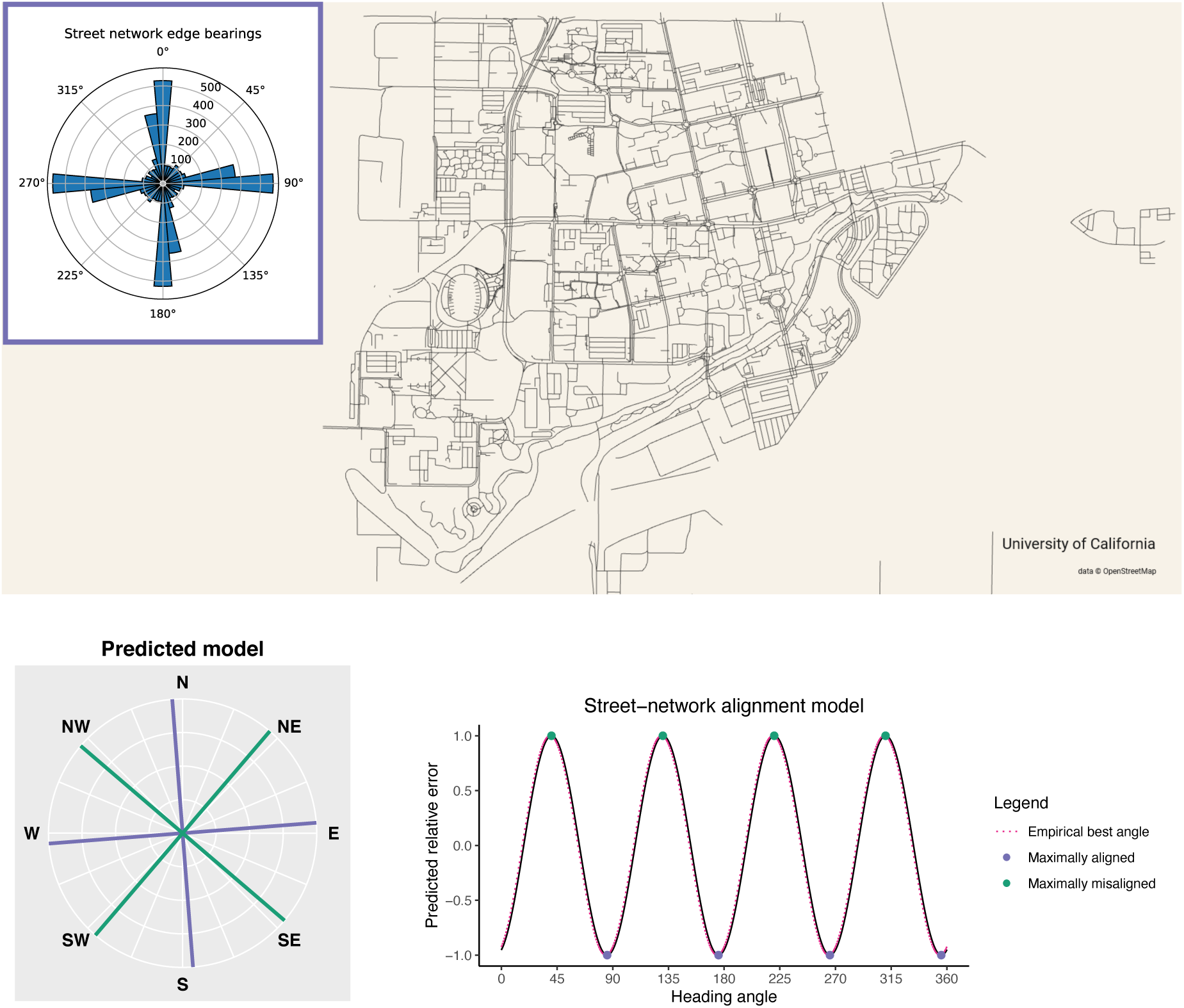
Participants’ long-term memories for their college campus exhibited a 4-fold alignment effect. Top: A map of the street network of the UC Davis campus; Inset: a polar histogram showing the distribution of bearings of the streets (in NSEW coordinates). Bottom panels: The predicted orientation of the 4-fold alignment effect for the street network bearings (i.e., the predicted model). Note, the preferred angle of the alignment model will be oriented toward the directions with the maximal error (i.e., the model has values of 1 at the maximal error and values of −1 at the minimal error); thus, we shifted the preferred alignment model by 45 degrees. Bottom right: Interestingly, the model uncovered the typical alignment of the campus, which is oriented slightly counterclockwise from NSEW, thus matching the street network edge bearings (i.e., the campus is rotated slightly counterclockwise from being the NSEW axes).

#### 3.2.3 Campus task: Evidence that the JRD Task, the Map Task, and the Distance Task recruit common underlying representations

We next tested whether the JRD Task, the Map Task, and the Distance Task recruit common underlying representations for the campus task. First, we observed a significant relationship between performance for all comparisons: 1) the JRD Task and the Map Task (Pearson’s *r* = −0.55, *t*_105_ = −6.77, *p =* 7.43×10^−10^, *BF*_10_ = 14890139; Spearman’s *ρ* = −0.64, *S* = 336438, *p <* 2.2×10^−16^; Kendall’s *τ* = −0.47, *z* = −7.13, *p* = 9.96 × 10^−13^), 2) the Distance Task and the Map Task (Pearson’s *r* = 0.52, *t*_102_ = 6.17, *p =* 1.42×10^−8^, *BF*_10_ = 9.7×10^5^; Spearman’s *ρ* = 0.49, *S* = 96110, *p <* 2.3×10^−7^; Kendall’s *τ* = −0.35, *z* = 5.31, *p* = 1.07 × 10^−7^), and 3) the JRD Task and the Distance Task (Pearson’s *r* = −0.51, *t*_102_ = −5.97, *p =* 3.53×10^−8^, *BF*_10_ = 4.17×10^5^; Spearman’s *ρ* = −0.57, *S* = 292472, *p <* 7.9×10^−10^; Kendall’s *τ* = −0.40, *z* = −6.02, *p* = 1.78 × 10^−9^; see Figure 4).

**Figure 4:**
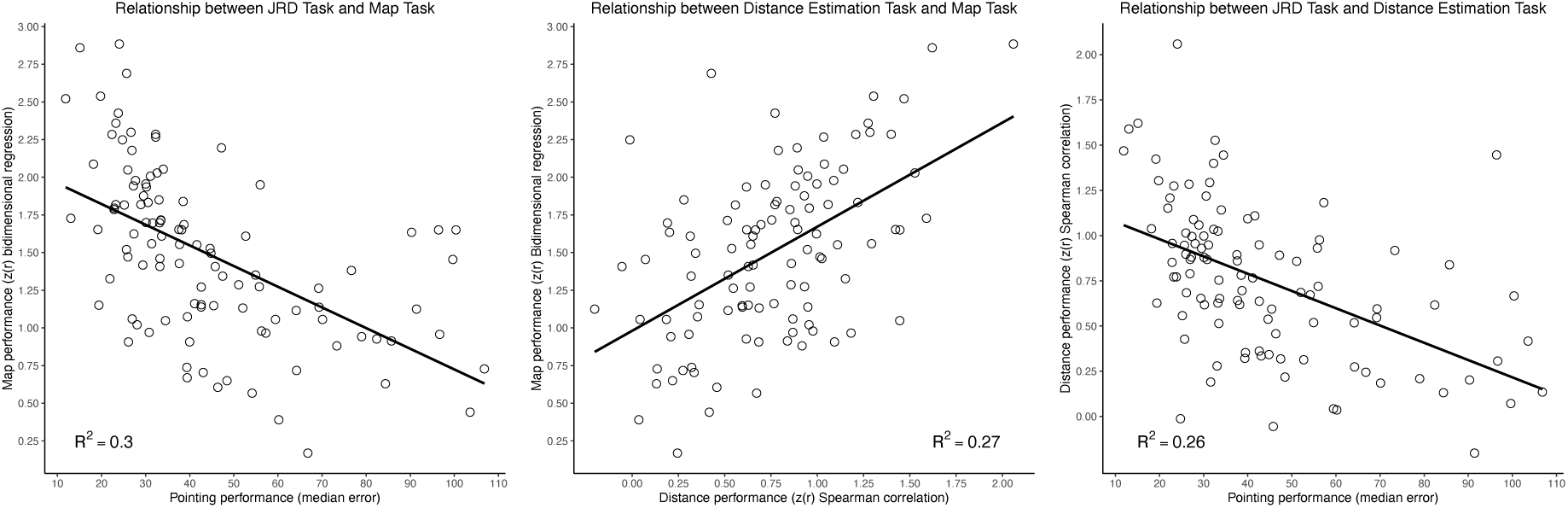
Evidence that participants recruit similar underlying cognitive representations to solve the JRD Task, the Map Task, and the Distance Estimation Task for their college campus. The correlation between performance was significant for the comparison between the JRD Task and the Map Task (see left panel), the Distance Estimation Task and the Map Task (see middle panel), and the JRD Task and the Distance Estimation Task (see right panel).

Second, we tested the hypothesis that error patterns would be correlated by employing a partial correlation approach. Here, we analyzed the partial correlations between 1) performance on the JRD Task and the Map Drawing Task, 2) the Distance Task and the Map Drawing Task (note that we could not compare performance between the JRD Task and Distance Task since these do not share a common format or straightforward comparison; see 2.5 and Appendix).

The Fisher’s r-to-z transformed circular correlation coefficients were significantly greater than zero for both the comparison of 1) the JRD Task and the Map Drawing Task (mean = 0.33, *t*_104_ = 9.97, *p <* 2.2×10^−16^, *BF*_10_ = 7.54 × 10^13^) and 2) the Distance Task and the Map Drawing Task (mean = 0.31, *t*_103_ = 11.78, *p <* 2.2×10^−16^, *BF*_10_ = 5.72×10^17^; see Figure 5).

**Figure 5:**
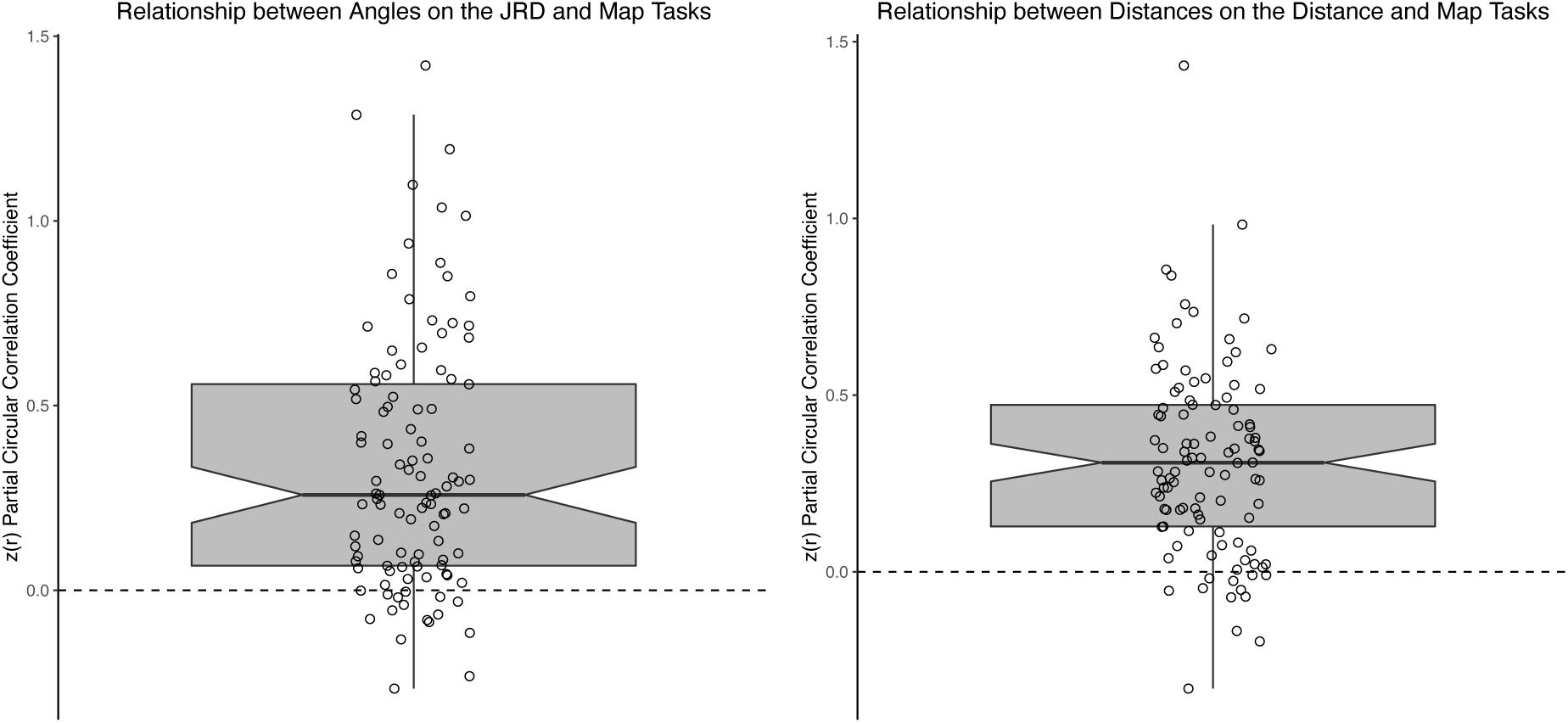
Evidence that participants recruit similar underlying cognitive representations to solve the Map Task as the JRD Task and the Distance Estimation Task for their college campus. Left panel: The Fisher’s r-to-z transformed partial circular correlation coefficients between responses on the Map Task and the JRD Task were significantly greater than zero (mean = 0.33, *t*_104_ = 9.97, *p <* 2.2×10^−16^, *BF*_10_ = 7.55×10^13^). Right panel: The Fisher’s r-to-z transformed partial circular correlation coefficients between responses on the Map Task and the Distance Estimation Task were significantly greater than zero (mean = 0.31, *t*_103_ = 11.78, *p <* 2.2×10^−16^, *BF*_10_ = 5.72×10^17^). Each dot indicates an individual participant and the boxplot depicts the middle 25th to 75th percentile of the data.

Taken together, our results replicate and extend upon our previous research using virtual environments (Huffman & Ekstrom, 2019) and provide evidence to suggest that the JRD Task, the Map Task, and the Distance Task recruit similar cognitive representations, even for real-world environments (note that we also added a new task here: The Distance Task; also see below for a replication within the Hometown data and the virtual environment data).

### 3.3 Discussion

The results from the real-world part of this experiment provide two sources of evidence to suggest that participants recruit cognitive heuristics to solve the tasks that result in violations of Euclidean geometry. First, the participants’ responses on the Map Task were better accounted for by an Affine model than a Euclidean model. These results suggest that the participants’ maps exhibit a significant degree of distortion, which is likely due to remembering landmarks as closer together, in some cases than they actually were. This could relate to the heuristic of “location is close to Y”. Second, we observed a significant alignment effect on pointing accuracy in the JRD task, and the angle of the alignment effect was related to the street bearings. These results suggest that participants used an alignment heuristic in their pointing responses, even for a large-scale, well-learned environment that is relevant to their daily lives (i.e., their college campus), for example, using north-south-east-west as a way to associate landmarks.

Additionally, our results provide evidence that participants recruit partially overlapping cognitive representations to solve the Map Task, the JRD Task, and the Distance Estimation Task for the real-world and virtual environments (see below for data from the virtual environment). We observed significant correlations in performance on these three tasks, thus suggesting that individual differences in spatial memory performance are correlated between these tasks. More directly, our partial correlation analysis allowed us to assess the relationship among the pattern of errors on these tasks, which provides a stronger test of the similarity of the cognitive representations that participants use to solve these tasks (see Huffman & Ekstrom, 2019). Here, we observed significant partial correlations between angular responses on the Map Task and the JRD Task as well as between the distance responses on the Map Task and the Distance Estimation Task, thus revealing a significant correlation among the pattern of errors on these tasks. Importantly, we observed these effects for both the real-world and the virtual environment (see below for data from the virtual environment), thus strongly suggesting that participants use partially overlapping underlying cognitive representations to solve these tasks. These results replicate and extend our previous work and highlight the construct validity of these tasks in tapping into the underlying cognitive representation of spatial memory.

## 4 Experiment 2

In Experiment 2, we tested whether real-world, large-scale, behaviorally relevant environments highly familiar but remotely learned spatial environments would show similar distortions. Here, we studied participants’ memory for their hometown. This provided a test to see whether highly learned environments, which presumably college-age students had extensive experience with growing up, would also show systematic distortions. The experiment began by asking participants to input the locations of 10 familiar landmarks in their hometown. We again ran the virtual environment task as well (in a separate session; see below for the results).

### 4.1 Methods

The design of Experiment 2 was similar to Experiment 1. However, in this task we tested participants’ knowledge of landmarks in their hometowns. Specifically, participants were asked to name (by typing) 10 landmarks that were most familiar to them from their hometowns. At the end of the hometown session, participants indicated the locations of the landmarks using Google’s “My Maps” (for more details see 2.3). Participants also completed two rounds of the Mental Rotation Task to assess how much small-scale spatial ability relates to large-scale spatial cognition.

We also included additional questionnaires, which we did not analyze here: a 5-point GPS use questionnaire (Ruginski et al., 2019) and we asked participants how many hours of video games they play on average per week, including ranking on a scale from 1 to 5 how often they play 2D vs. 3D video games (e.g., Clemenson et al., 2019; Clemenson & Stark, 2015).

### 4.2 Results

#### 4.2.1 Hometown task: Evidence that participants’ memories for their hometown are better explained by distorted representations than by Euclidean maps

As in our analysis of participants’ maps from the local university campus, we investigated whether participants’ maps were systematically distorted (i.e., better fit by an affine versus a Euclidean transformation; positive ΔAIC are in favor of the Euclidean model while negative values are in favor of the affine model). If participants formed metric, Euclidean maps of their hometowns (as might be expected by their long-term exposure to these environments) then we should observe evidence in favor of the Euclidean model. Conversely, if participants rely on heuristics and form distorted memories of their hometowns, then we should observe evidence in favor of the affine model. As can be seen in Figure 6, the ΔAIC was negative for the majority of our participants (the ΔAIC was less than 0 for 77.3% and less than −2 for 68.0% of our participants). In contrast, the Euclidean model provided a better fit (i.e., ΔAIC *>* 2) in a small percentage of participants (14.4%) and the models were inconclusive (i.e., −2 *<* ΔAIC *<* 2) in a small percentage of participants (17.5%). Two-tailed Wilcoxon signed rank tests revealed that the median ΔAIC, −5.99, was significantly less than 0 (*p* = 8.28 × 10^−14^) and significantly less than −2 (*p* = 7.64 × 10^−8^; note: as mentioned above, |ΔAIC| *>* 2 is typically taken to as evidence of a significant difference between models; Nakaya, 1997). Thus, the majority of our participants’ map data are better explained by the affine model relative to the Euclidean model. These results replicate and extend our findings from the local university campus and suggest that the structure of participants’ knowledge of their hometowns is better explained by distorted, heuristic representations than by metric, Euclidean maps (e.g., Nakaya, 1997).

**Figure 6:**
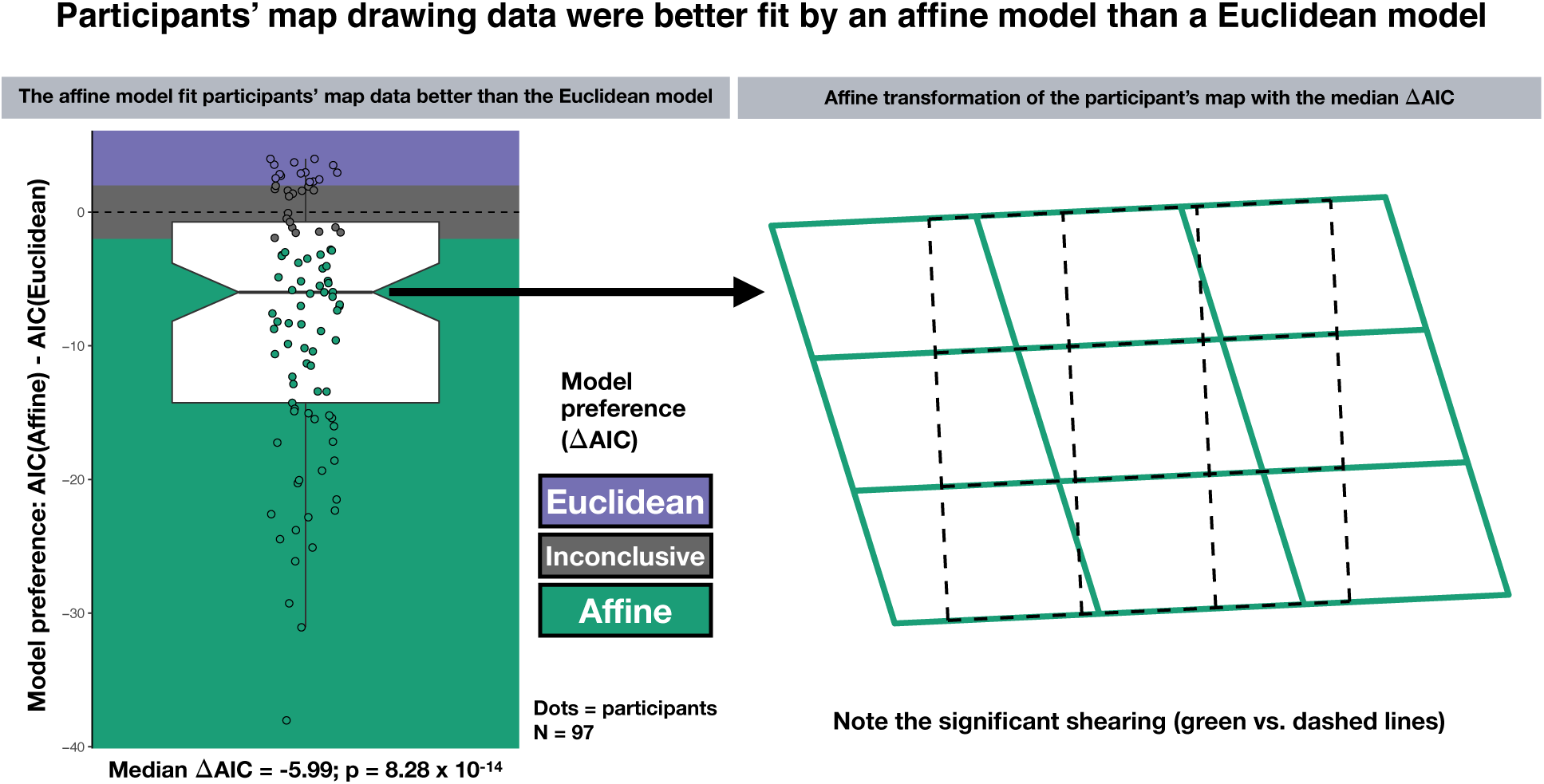
Evidence that participants’ memories for their hometown are distorted and do not follow Euclidean geometry. Each participant’s map drawing data were fit with Euclidean and affine models. This figure depicts the ΔAIC between the two model fits, where negative values are in favor of the affine model and positive values favor the Euclidean model. The ΔAIC for the majority of participants was less than −2, which is typically taken to be significant evidence in favor of the model. Moreover, a Wilcoxon signed rank test revealed that the median ΔAIC, −5.99, was significantly less than 0 (*p* = 8.28 × 10^−14^). Each dot indicates an individual participant and the boxplot depicts the middle 25th to 75th percentile of the data. The colors indicate model preference, where the Euclidean model is preferred if ΔAIC *>* 2, the affine model is preferred if ΔAIC *<* −2, and the model comparison is inconclusive if −2 *<* ΔAIC *<* 2.

As an exploratory analysis, we next investigated whether distortions in participants’ maps could be at least partially attributable to differences in the nature of the environment in which they grew up. For example, recent research found that people that grow up in less grid-like cities tend to perform better on new spatial learning (Coutrot et al., 2022). Thus, here, we explore whether we would observe a similar effect in the participants’ ability to draw maps of their hometown. Specifically, based on the previous literature, we predicted that we would find that participants that grew up in more grid-like cities (Coutrot et al., 2022).

would exhibit more evidence in favor of distorted representations (i.e., affine model) relative to participants that grew up in less grid-like cities. Thus, we calculated the street network entropy (a measure of how ordered the streets are; lower values indicate more grid-like cities whereas higher values indicate less grid-like cities) and compared it to the model preference (i.e., the Δ*AIC* for the affine model – the Euclidean model). We found a positive correlation between street network entropy and the model preference (Δ*AIC* for the affine model – the Euclidean model; Pearson r = 0.286, t_95_ = 0.0044, BF_10_ = 10.8; Spearman rho = 0.265, p = 0.0086; see Figure 7), thus suggesting that participants that grew up in more grid-like cities exhibited greater degrees of distortions in memory (i.e., model preference for the affine model vs. the Euclidean model). Altogether, these exploratory results provide new evidence to suggest that part of that distortions in participants’ memories for their hometown can be partially explained by the nature of the environment itself—i.e., how grid-like the environment is—which suggests that there could be developmental and practice differences as a function of the environments in which we grow up, thus building on previous research in memory (i.e., model preference for the affine model vs. the Euclidean model).

**Figure 7:**
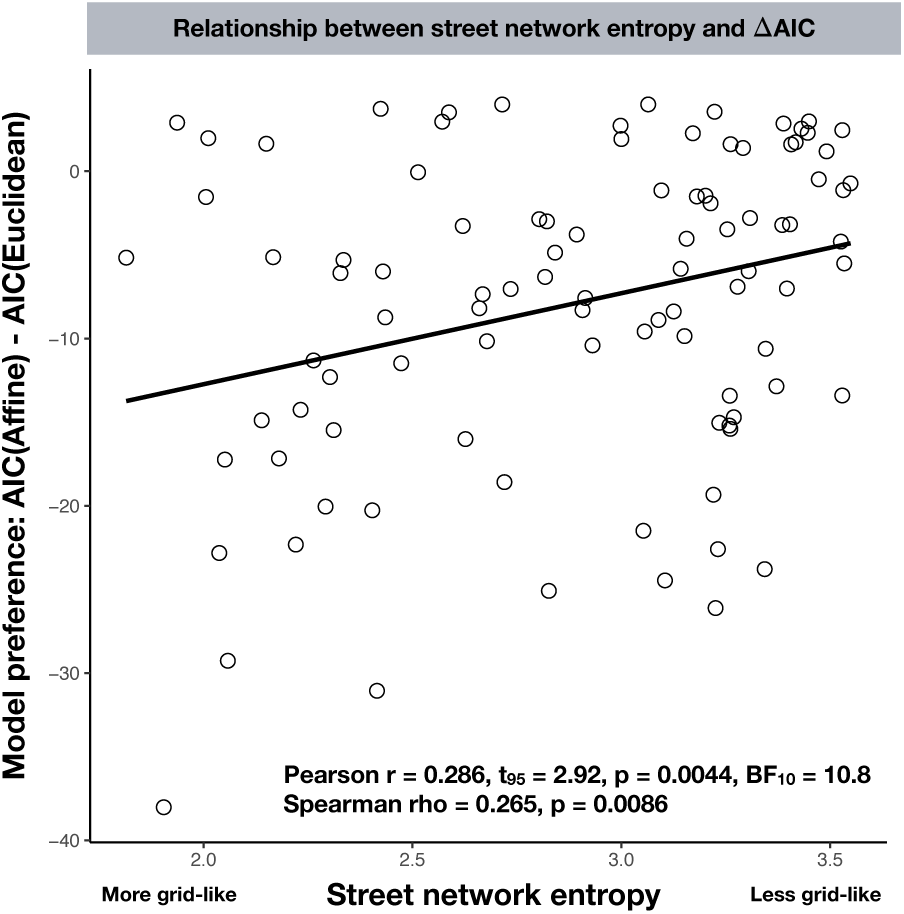
In an exploratory analysis, we observed a significant positive correlation between street network entropy and the model preference (Δ*AIC* for the affine model – the Euclidean model) for participants’ maps for their hometown. This suggests that participants that grew up in more grid-like-cities exhibited greater degrees of distortions.

We next tested the hypothesis that participants’ (very) long-term spatial representations for their hometown cities would exhibit alignment effects. We used a within-participant cross-validation approach to account for the fact that each participant had a different hometown and thus, would potentially exhibit a different orientation of alignment effects. We found a significant cross-validated 4-fold alignment effect in pointing errors (mean *β* = 0.077, *t*_96_ = 3.72, *p* = 0.00033, *BF*_10_ = 62.1; see Figure 8). These results provide additional evidence for systematic distortions in memory for very well learned environments. Specifically, participants’ memories appear to exhibit an alignment heuristic, thus suggesting that boundaries likely explain some of the systematic distortions in memory, which dovetails nicely with the findings that participants’ maps were better fit by the affine model than the Euclidean model.

**Figure 8:**
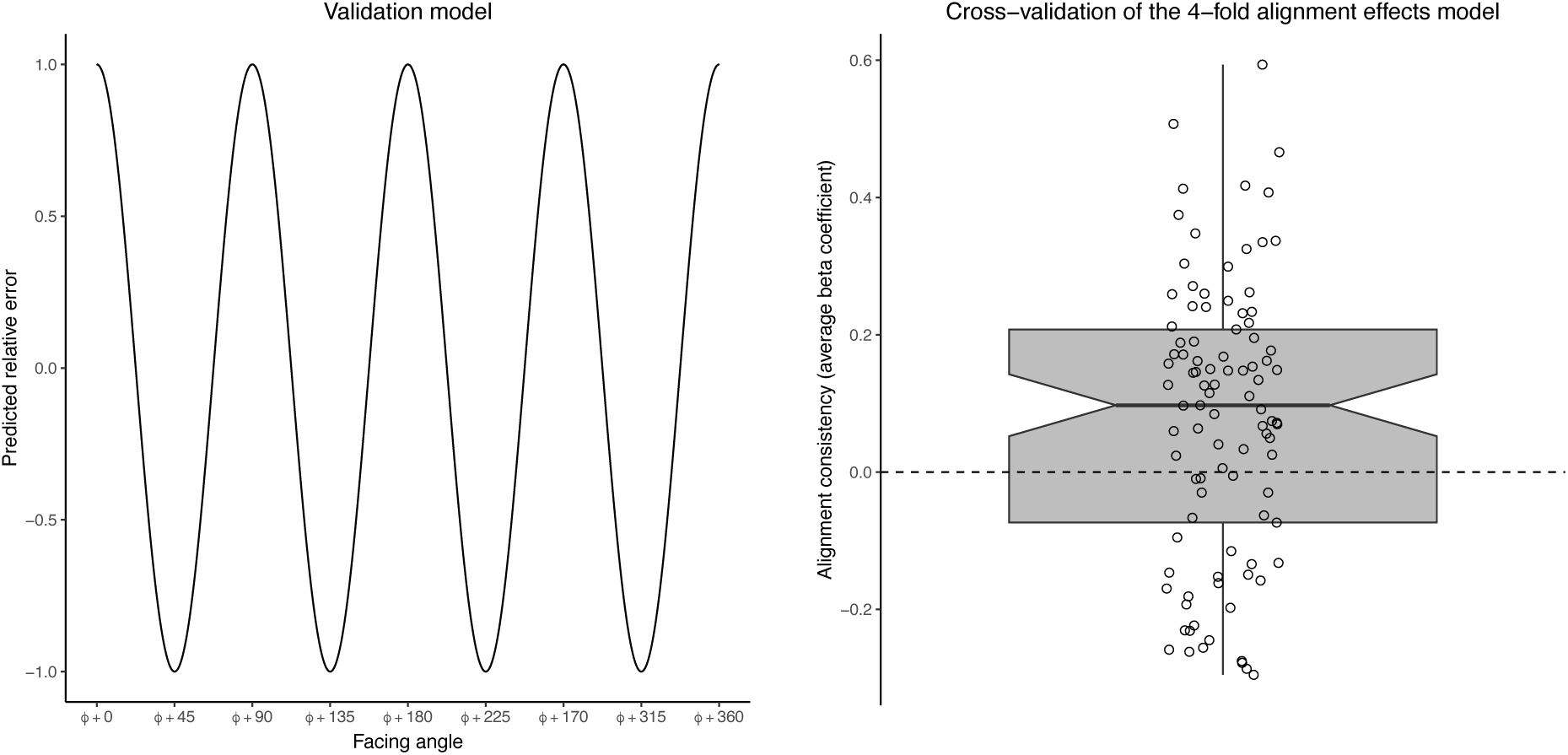
Participants’ long-term memories for their hometown cities exhibited a 4-fold alignment effect. Left: We used a within-subject cross-validation approach in which we trained an alignment effects model on a subset of data and tested the consistency of its effect using a validation model (*ϕ* = preferred orientation from the training data; the figure shows a compass rose of the validation model). Right: We found a significant cross-validated 4-fold alignment effect.

#### 4.2.2 Hometown Task: Evidence that the JRD Task, the Map Task, and the Distance Estimation Task Recruit Common Underlying Representations

We next tested whether the JRD Task, the Map Task, and the Distance Estimation Task recruit common underlying representations for the hometown task. First, we tested the correlations between performance across these three tasks. We observed a significant relationship between performance for all three comparisons: 1) the JRD Task and the Map Task (Pearson’s *r* = −0.27, *t*_95_ = −2*.7*4, *p* = .0073, *BF*_10_ = 7.07; Spearman’s *ρ* = −0.36, *S* = 206638, *p =* .00034; Kendall’s *τ* = −0.25, *z* = −3.64, *p* = .00027), 2) the Distance Estimation Task and the Map Task (Pearson’s *r* = 0.44, *t*_92_ = 4.71, *p =* 8.86.×10^−6^, *BF*_10_ = 2655.5; Spearman’s *ρ* = 0.40, *S* = 82458, *p <* 6.15×10^−5^; Kendall’s *τ* = −0.28, *z* = 3.98, *p* = 7.04 × 10^−5^), and 3) the JRD Task and the Distance Estimation Task (Pearson’s *r* = −0.34, *t*_92_ = −3.44, *p* = .00088, *BF*_10_ = 44.3; Spearman’s *ρ* = −0.33, *S* = 183800, *p* < .0013; Kendall’s *τ* = −0.23, *z* = −3.32, *p* = .00089; see Figure 9).

**Figure 9:**
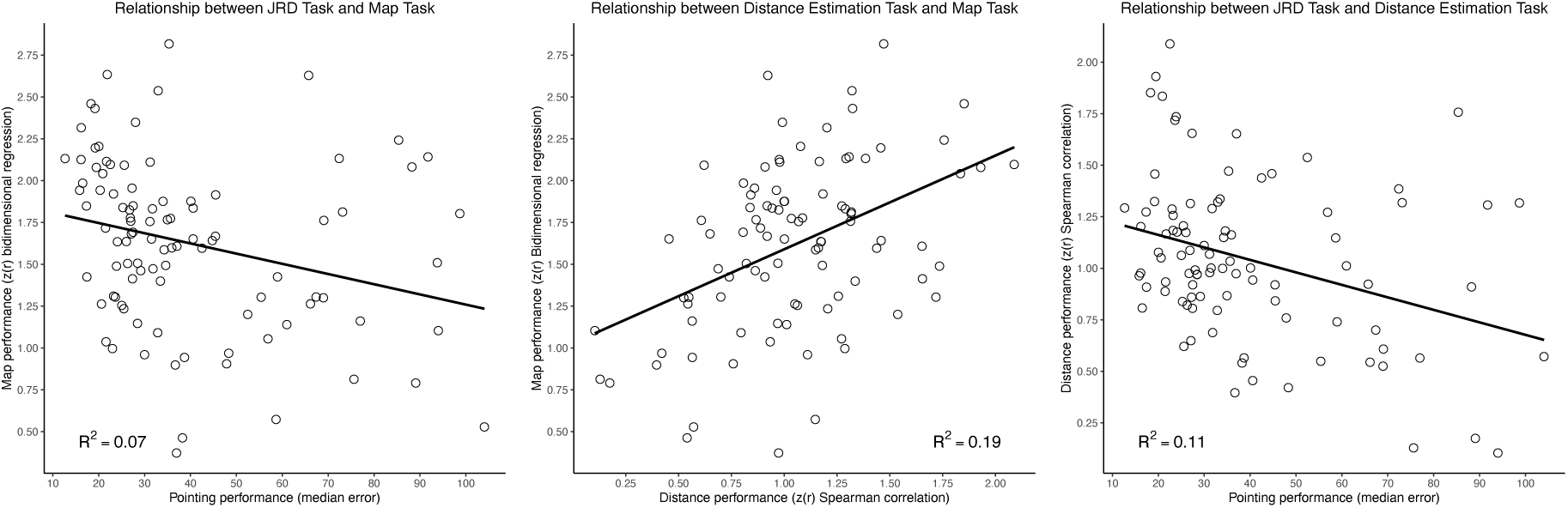
Evidence that participants recruit similar underlying cognitive representations to solve the JRD Task, the Map Task, and the Distance Estimation Task for their college campus. The correlation between performance was significant for the comparison between the JRD Task and the Map Task (see left panel), the Distance Estimation Task and the Map Task (see middle panel), and the JRD Task and the Distance Estimation Task (see right panel).

Second, the Fisher’s r-to-z transformed partial circular correlation coefficients were significantly greater than zero for the JRD Task and the Map Task (mean = 0.36, *t*_96_ = 9.62, *p* = 9.78×10^−16^, *BF*_10_ = 6.57 × 10^12^) and for the Distance Estimation Task and the Map Task (mean = 0.32, *t*_93_ = 10.45, *p <* 2.2 × 10^−16^, *BF*_10_ = 2.55 × 10^14^; see Figure 10). It is important to note that our partial correlation analysis measured the relationship of the pattern of responses between the two tasks while partialing out the effect of the answers; thus, these results are indicative of similarity between participants’ responses on the two tasks above and beyond what would be predicted from a mutual correlation with the answers (for more information about our partial circular correlation technique please see 2.5 and Appendix). The results of the partial correlation analysis suggest that participants recruit similar underlying cognitive representations to solve the JRD Task, the Map Task, and the Distance Estimation Task. Therefore, these results extend our current and previous findings from virtual environments (also see Huffman & Ekstrom, 2019) to real-world environments, including participants’ hometowns, as well as a new task: Distance Estimation Task, and (at least partially) mitigate against the concern that there is something artificial about our virtual navigation task.

**Figure 10:**
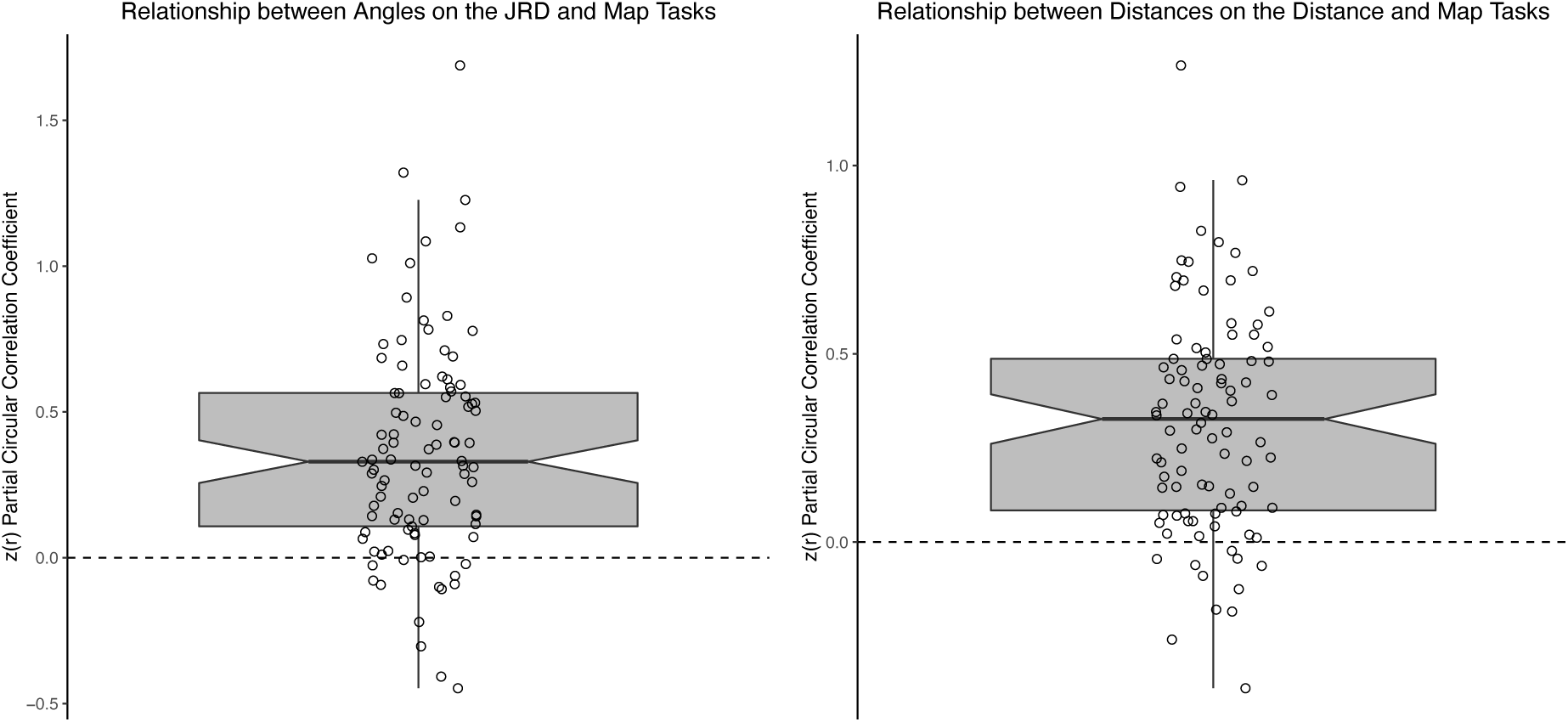
Evidence that participants recruit similar underlying cognitive representations to solve the Map Task as the JRD Task and the Distance Estimation Task for their hometowns. Left panel: The Fisher’s r-to-z transformed partial circular correlation coefficients between responses on the Map Task and the JRD Task were significantly greater than zero (mean = 0.36, *t*_96_ = 9.62, *p* = 9.78×10^−16^, *BF*_10_ = 6.57×10^12^). Right panel: The Fisher’s r-to-z transformed partial circular correlation coefficients between responses on the Map Task and the Distance Estimation Task were significantly greater than zero (mean = 0.32, *t*_93_ = 10.45, *p <* 2.2 × 10^−16^, *BF*_10_ = 2.55 × 10^14^). Each dot indicates an individual participant and the boxplot depicts the middle 25th to 75th percentile of the data.

### 4.3 Discussion

The findings of Experiment 2 further support the idea that people use heuristics and distorted underlying representations to solve spatial memory tasks, even when they are tested on an environment that is extremely well learned, like their hometown. Participant responses on the Map Task exhibited systematic distortions, as evidenced by an affine model fitting their responses better than the Euclidean model. Importantly, the effects were remarkably similar for both the UC Davis campus (Experiment 1) and the hometown (Experiment 2) environments, thus suggesting the underlying representations across these two timescales are similar. In addition, we observed a 4-fold alignment effect for the pointing errors made in the JRD task, suggesting that people relied on alignment heuristics to complete their pointing judgments. Not only do participants have distorted maps, but they also rely on heuristics when asked to point between locations that they have known their entire lives.

We also replicated the findings of Experiment 1 regarding the relationship between performance on each of our spatial memory tasks, providing more evidence participants use similar underlying cognitive representations during different tests of their memory of the environment. Within the hometown environment, our partial correlation analysis revealed a significant relationship between performance on the JRD, Distance Estimation, and Map Tasks. This finding extends that of Experiment 1 by strengthening the existing evidence, this time when probing participants about an extremely well-learned, real-world environment, that individual differences are consistent across these three spatial memory tasks. In addition, we replicated previous findings that repeated navigation increased performance, although we did observe a large spread in performance throughout the JRD task. These results demonstrate a wide array of individual differences in spatial memory tasks, regardless of how well-learned the environment is, and these differences remain consistent across tasks.

## 5 Experiment 1, Novel Virtual Environment

### 5.1 Method

Here, all participants from Experiment 1 also navigated within a novel virtual environment. Please see more details above in the General Methods section.

### 5.2 Results

#### 5.2.1 Novel Virtual Environment: Evidence of a Boundary Expansion Effect

We next tested the hypothesis that the map-drawing performance for the virtual environment would be subjected to systematic distortions. We found evidence that participants drew the locations of landmarks that were proximal to the boundaries as being closer to the boundaries than they actually were; an effect that we refer to as a “boundary expansion effect.” Specifically, during the Map-Drawing Task, the participants saw an overhead view of the environment in which the walls were still in visible. We found that participants made a key distortion in their memory such that they tended to draw the coordinates closer to the wall than they actually were, which we term the “boundary expansion effect” (e.g., for the raw coordinates see Figure 11). Moreover, the bidimensional regression model exhibited a significant scaling effect, such that participant’s maps were significantly expanded relative to the actual maps (mean scaling factor = 0.79, *t*_123_ = −11.2, *p <* 2.2 × 10^16^; BF_10_ = 3.41 × 10^17^; see Figure 11 and Figure 12).

**Figure 11:**
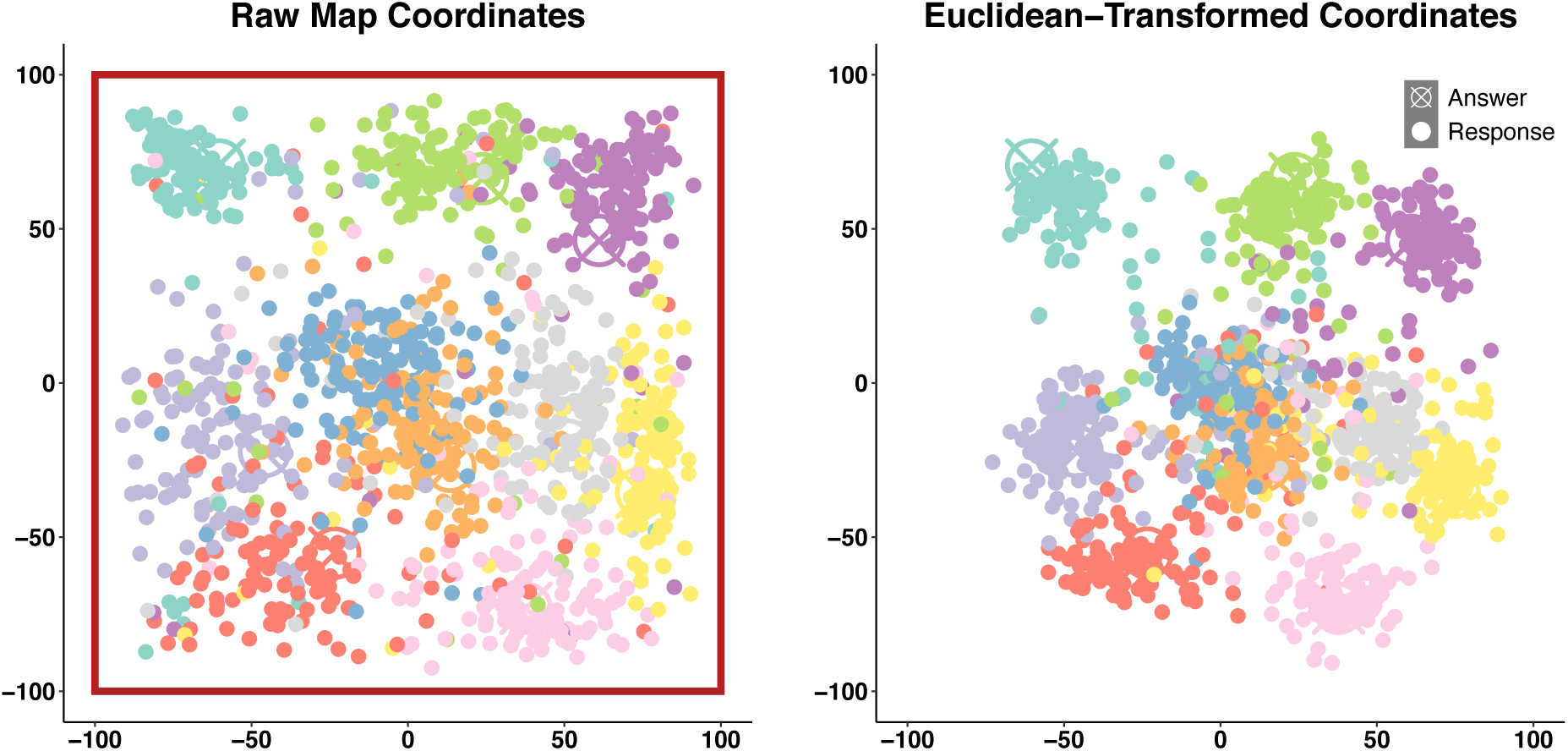
The participants’ maps were overall relatively accurate but displayed notable distortions. Left: The participants’ raw map responses (note: the red box depicts the locations of the boundaries of the environment; here, the participants drew maps that were significantly pulled toward the boundaries, termed a “boundary expansion effect”; also see Figure 12). Right: The Euclidean-transformed response coordinates. The circled x’s indicate the ground truth coordinates while the dots indicate individual participant coordinates and each location is depicted in a unique color.

**Figure 12:**
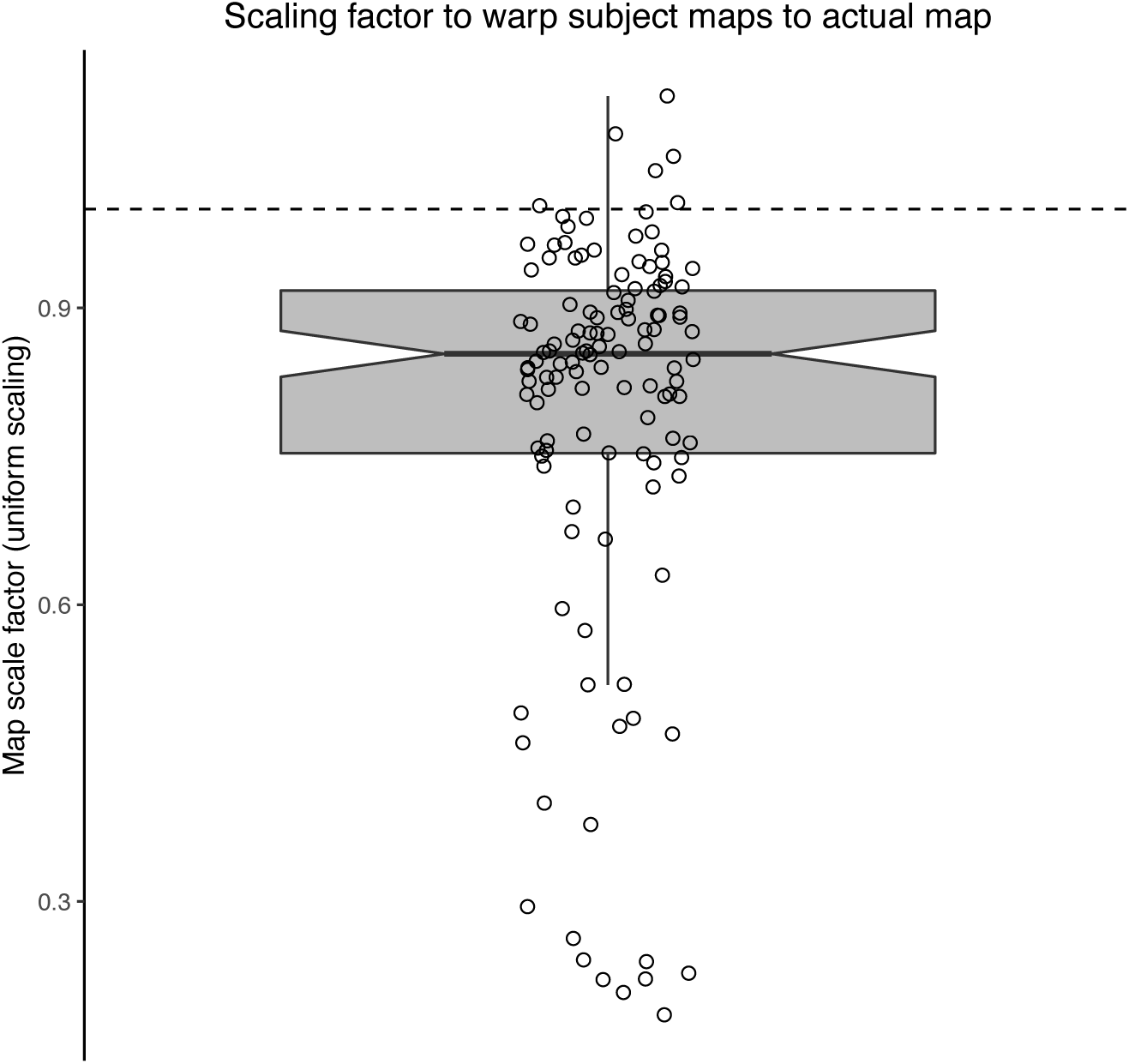
Evidence that participants warp their maps in a boundary-expansion effect (i.e., closer to the walls than the actual maps). Here, each dot indicates the (uniform) scaling factor required in the transformation from the participant map to the actual map. Each dot indicates the scale factor for an individual participant and the boxplot depicts the middle 25^th^ to 75^th^ percentile of the data.

Moreover, we found that, as a result of the boundary expansion effect, participants also placed the items that were near the boundary as being more aligned than they actually were, which is more evidence of a boundary-alignment heuristic underlying their memory performance (see Supplemental Material and Supplemental Figure 3; note that we used the bpnreg package for this analysis: Cremers & Klugkist, 2018).

#### 5.2.2 Novel Virtual Environment: Evidence for Alignment Effects in Spatial Memory

We next aimed to determine whether participants exhibited boundary alignment effects in their memory for the virtual environment (see Figure 1B). Specifically, we tested whether the imagined heading on the JRD trials would influence pointing performance such that participants would point more accurately on trials in which the imagined heading was relatively aligned to the main axes of the boundaries vs. mis-aligned (using a sinusoidal regression model). Consistent with our predictions, we found a significant effect of the alignment model parameter (alignment beta = 0.91, t = 5.68, p = 1.39 × 10^-8^; see Figure 13; note that the z-scored answer model parameter was also significant: answer beta = 0.16, t = 15.7, p = 2.0 × 10^-22^), thus providing evidence for a boundary-alignment effect.

**Figure 13:**
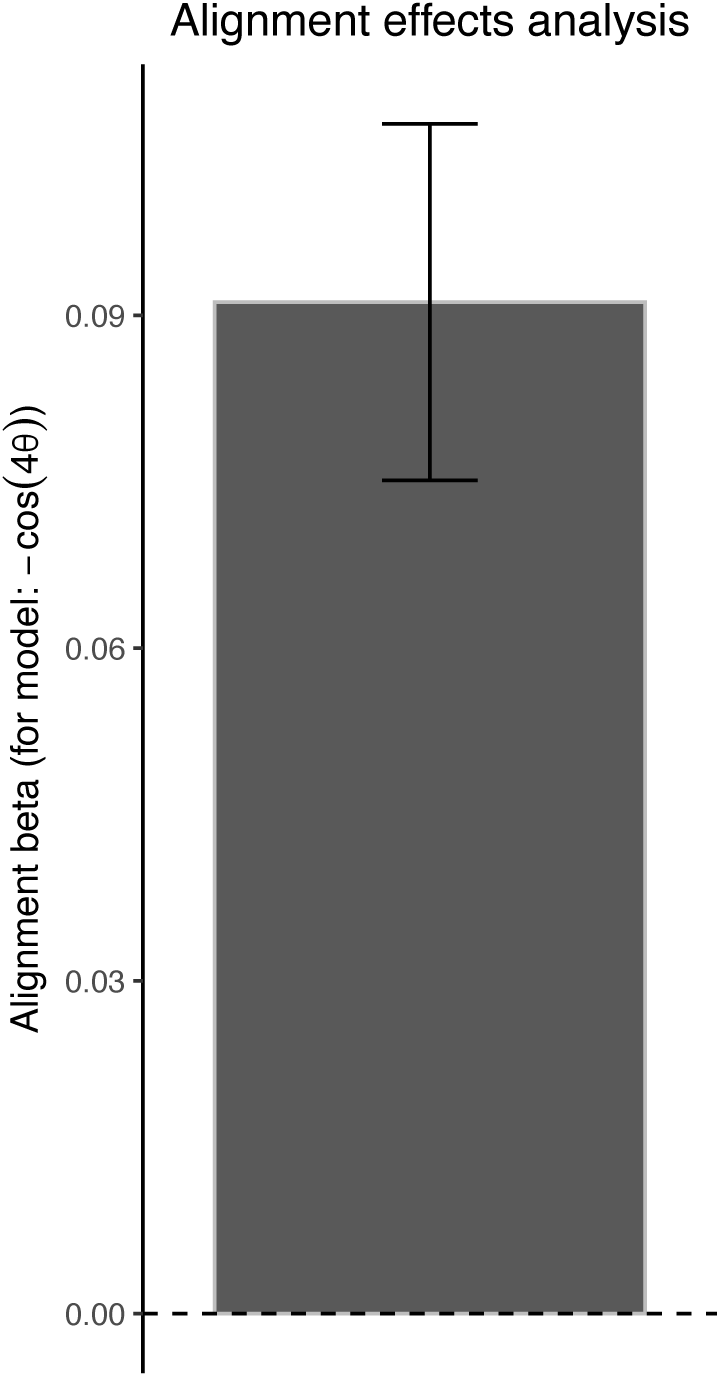
Evidence for an alignment effect within the JRD pointing accuracy (within the Experiment 1 data). The bar indicates the beta coefficient and the error bars indicate the standard errors from our mixed effects regression analysis.

#### 5.2.3 Novel Virtual Environment: Evidence that the JRD Task and the Map Task Recruit Common Underlying Representations

If participants recruit similar underlying cognitive representations to solve the JRD Task and the Map Task and both show similar patterns of systematic distortions, then we should observe a significant relationship between performance on the two tasks and we should also observe evidence of similar pattern of errors in responses within an individual.

Our results provide evidence for both of these effects. First, we observed a significant relationship between performance on the JRD Task and the Map Task (Pearson’s *r* = −0.74, *t*_122_ = −12.18, *p <* 2.2×10^−16^, *BF*_10_ = 2.60×10^19^; Spearman’s *ρ* = −0.61, *S* = 511724, *p <* 2.2×10^−16^; Kendall’s *τ* = −0.44, *z* = −7.19, *p* = 6.18 × 10^−13^; see Figure 14).

**Figure 14:**
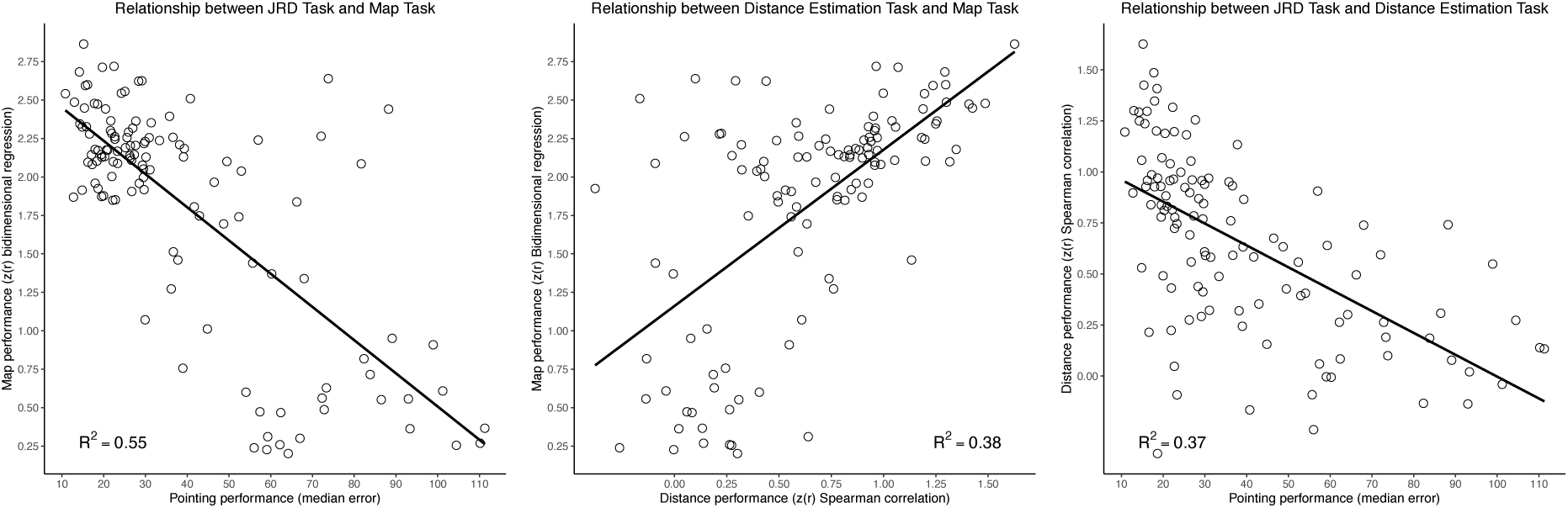
Relationship between the Map Task, JRD Task, and Distance Estimation Task for the virtual environment. Left: The relationship between performance on the JRD Task and the Map Task. Center: The relationship between performance on the Distance Estimation Task and the Map Task. Right: The relationship between performance on the JRD Task and the Distance Estimation Task.

Second, we tested the hypothesis that error patterns would be correlated by employing a partial correlation approach (see 2.5 and Appendix). The Fisher’s r-to-z transformed circular correlation coefficients were significantly greater than zero (mean = 0.21, *t*_123_ = 9.74, *p <* 2.2×10^−16^, *BF*_10_ = 1.01 × 10^14^; see Figure 15). Taken together, these results replicate our previous research (Huffman & Ekstrom, 2019) and provide further evidence to suggest that these tasks recruit similar cognitive representations.

**Figure 15:**
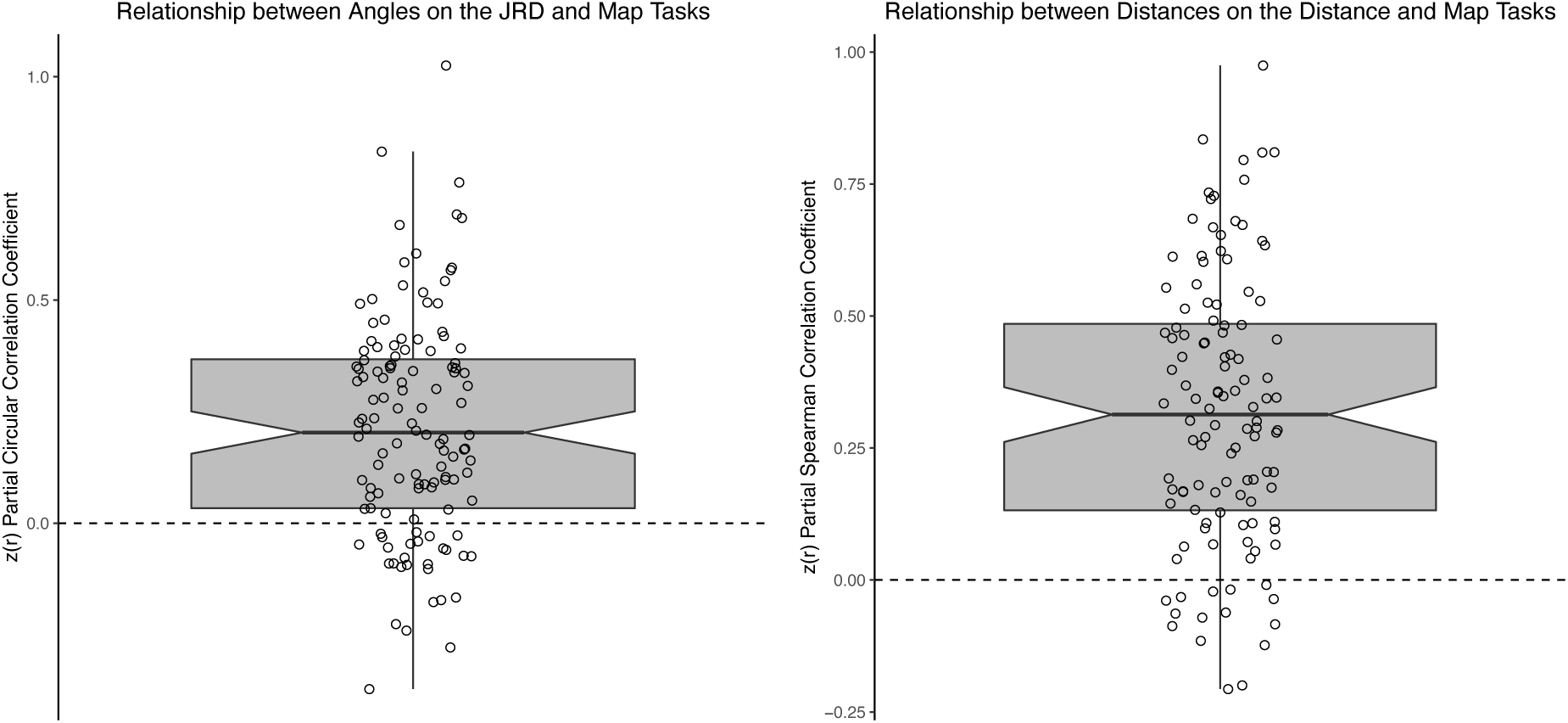
Evidence that participants recruit similar underlying cognitive representations to solve the Map Task as the JRD Task and the Distance Estimation Task for the virtual environment. Left panel: The Fisher’s r-to-z transformed partial circular correlation coefficients between the Map Drawing Task and the JRD Task were significantly greater than zero (mean = 0.21, *t*_123_ = 9.74, *p <* 2.2×10^−16^, *BF*_10_ = 1.01×10^14^). Right panel: The Fisher’s r-to-z transformed partial Spearman correlation coefficients between the Map Drawing Task and the JRD Task were significantly greater than zero (mean = 0.32, *t*_115_ = 13.44, *p <* 2.2×10^−16^, *BF*_10_ = 1.64×10^22^). Each dot indicates an individual participant and the boxplot depicts the middle 25th to 75th percentile of the data.

#### 5.2.4 Virtual environment: Evidence that the Distance Estimation Task and the Map Task recruit common underlying representations

Similar to our approach for the JRD Task and the Map Task, we employed two analyses to investigate the relationship between the Distance Estimation Task and the Map Task. First, we observed a significant relationship between performance on the two tasks (Pearson’s *r* = −0.61, *t*_115_ = −8.26, *p <* 2.81×10^−13^, *BF*_10_ = 2.42×10^10^; Spearman’s *ρ* = −0.67, *S* = 446,108, *p <* 2.2×10^−16^; Kendall’s *τ* = −0.49, *z* = −7.87, *p* = 3.47 × 10^−15^; see Figure 14). Second, the Fisher’s r-to-z transformed partial Spearman rank correlation coefficients between distance estimates on the Distance Estimation Task and the Map Task were significantly greater than zero when partialing out the effect of the answers (mean = 0.32, *t*_115_ = 13.44, *p <* 2.2 × 10^−16^, *BF*_10_ = 1.64 × 10^22^; see Figure 15). These results suggest that participants recruit similar representations to solve these tasks.

#### 5.2.4 Novel virtual environment: Evidence that the JRD Task and the Distance Estimation Task recruit common underlying representations

We observed a significant relationship between performance on the JRD Task and the Distance Estimation Task (Pearson’s r = −0.61, t_116_ = −8.29, p = 2.23 × 10^−13^, BF_10_ = 3.03 × 10^10^; Spearman’s ρ = −0.67, S = 457726, p < 2.20 × 10^−16^; Kendall’s τ = −0.49, z = −7.93, p = 2.13 × 10^−15^; see Figure 14).

#### 5.2.5 Novel virtual environment

Evidence that learning is enhanced by repeated navigation Our results reveal that pointing performance and confidence on the JRD Task improved with repeated rounds of navigation in the virtual environment (see Supplemental Information and Supplemental Figure 1), thus replicating our previous work (Huffman & Ekstrom, 2019).

#### 5.2.6 Comparison between performance on the real-world (campus) and virtual environment tasks

We next tested the correlations between performance on the real-world and virtual environment tasks. We found that the correlation between all three tasks was significant: JRD Task (Pearson’s *r* = 0.61, *t*_105_ = 7.92, *p* = 2.58 × 10^-12^, *BF*_10_ = 2872540527; Spearman’s *ρ* = 0.51, *S* = 100312, *p* = 3.55 × 10^-8^; Kendall’s *τ* = 0.38, *z* = 5.74, *p* = 9.36 × 10^-9^), Map Drawing Task (Pearson’s *r* = 0.33, *t*_105_ = 3.58, *p* = .00051, *BF*_10_ = 68.7; Spearman’s *ρ* = 0.31, *S* =140400, *p* = .0011; Kendall’s *τ* = 0.21, *z* = 3.22, *p* = .0012), and the Distance Estimation Task (Pearson’s *r* = 0.49, *t*_99_ = 5.54, *p* = 2.51 × 10^-7^, *BF*_10_ = 68194.74; Spearman’s *ρ* = 0.48, *S* =89226, *p* = 5.25 × 10^-7^; Kendall’s *τ* = 0.34, *z* = 5.09, *p* = 3.51 × 10^-7^; see Figure 16). Altogether, these correlations show that there is certainly shared variance between performance on the real-world and virtual environments; however, there is also a fair degree of unique variance (i.e., all of the r^2^’s < 0.4, thus explaining less than 40% of the variance), thus suggesting there may also be unique aspects of the real-world navigation task that cannot be fully captured with the novel virtual environment.

**Figure 16:**
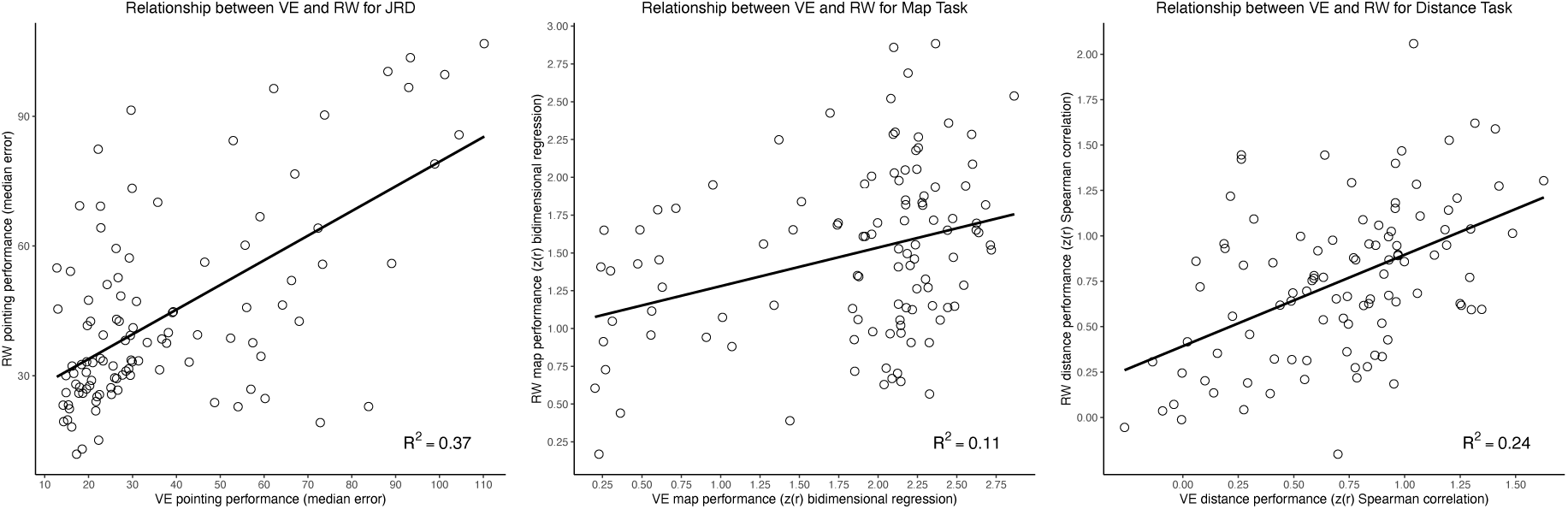
The correlations were significant for the comparison between performance on the virtual environment and the real world (campus) for the JRD Task (see left panel), the Map Task (see middle panel), and the Distance Estimation Task (see right panel).

### 5.3 Discussion

There are several key sets of findings in the present data. The first set of findings relate to the map drawing and JRD pointing responses, which both provide evidence of alignment effects. First, participants drew maps that were warped toward the boundaries of the environment, termed the “boundary expansion effect.” Moreover, the “boundary expansion effect” also caused participants to draw outer landmarks as more aligned to the external boundaries than they actually were (i.e., a form of a boundary alignment effect). Importantly, the fact that we included the walls of the environment during the map drawing task allowed us to do tjos more detailed analysis of the maps. Additionally, as in the real-world data, we observed significant boundary alignment effects in JRD responses.

We also observed robust evidence for correlations between performance across measures. As in the real-world data, we observed robust correlations between performance on the JRD Task, the Map Drawing Task, and the Distance Estimation Task. Moreover, we again observed significantly correlated errors between the JRD Task and the Map Drawing Task and the Distance Estimation Task and the Map Drawing Task.

Altogether, the data from the Map Drawing Task and the JRD Task further elucidates the nature of spatial memory distortions by suggesting that alignment effects are a prominent heuristic used across tasks. Moreover, the strong relationship between tasks bolsters both the notion that participants use partially overlapping representations to solve the task and that such representations may be similarly distorted. We return to these issues in the Discussion for Experiment 2, in which we replicate these findings, as well as in the General Discussion.

## 6 Experiment 2, Novel Virtual Environment (Replication with an independent data set within the same environment)

### 6.1 Method

Here, the methods were identical to those in Experiment 1, thus our approach here allows a direct replication of our findings in the previous section.

### 6.2 Results

#### 6.2.1 Novel Virtual Environment: Evidence of a Boundary Expansion Effect

As in Experiment 1, we found evidence that participants drew the locations of landmarks that were proximal to the boundaries as being closer to the boundaries than they actually were, thus we replicated the “boundary expansion effect.” Specifically, during the Map-Drawing Task, the participants saw an overhead view of the environment in which the walls were still in visible. We found that participants made a key distortion in their memory such that they tended to draw the coordinates closer to the wall than they actually were, which we term the “boundary expansion effect” (e.g., for the raw coordinates see Figure 17). Moreover, the bidimensional regression model exhibited a significant scaling effect, such that participant’s maps were significantly expanded relative to the actual maps (mean scaling factor = 0.81, *t*_114_ = −10.6, *p <* 2.2 × 10^16^; BF_10_ = 4.86 × 10^15^; see Figure 17 and Figure 18). Moreover, we found that, as a result of the boundary expansion effect, participants also placed the items that were near the boundary as being more aligned than they actually were, which is more evidence of a boundary-alignment heuristic underlying their memory performance (see Supplemental Material and Supplemental Figure 3; note that we used the bpnreg package for this analysis: Cremers & Klugkist, 2018).

**Figure 17:**
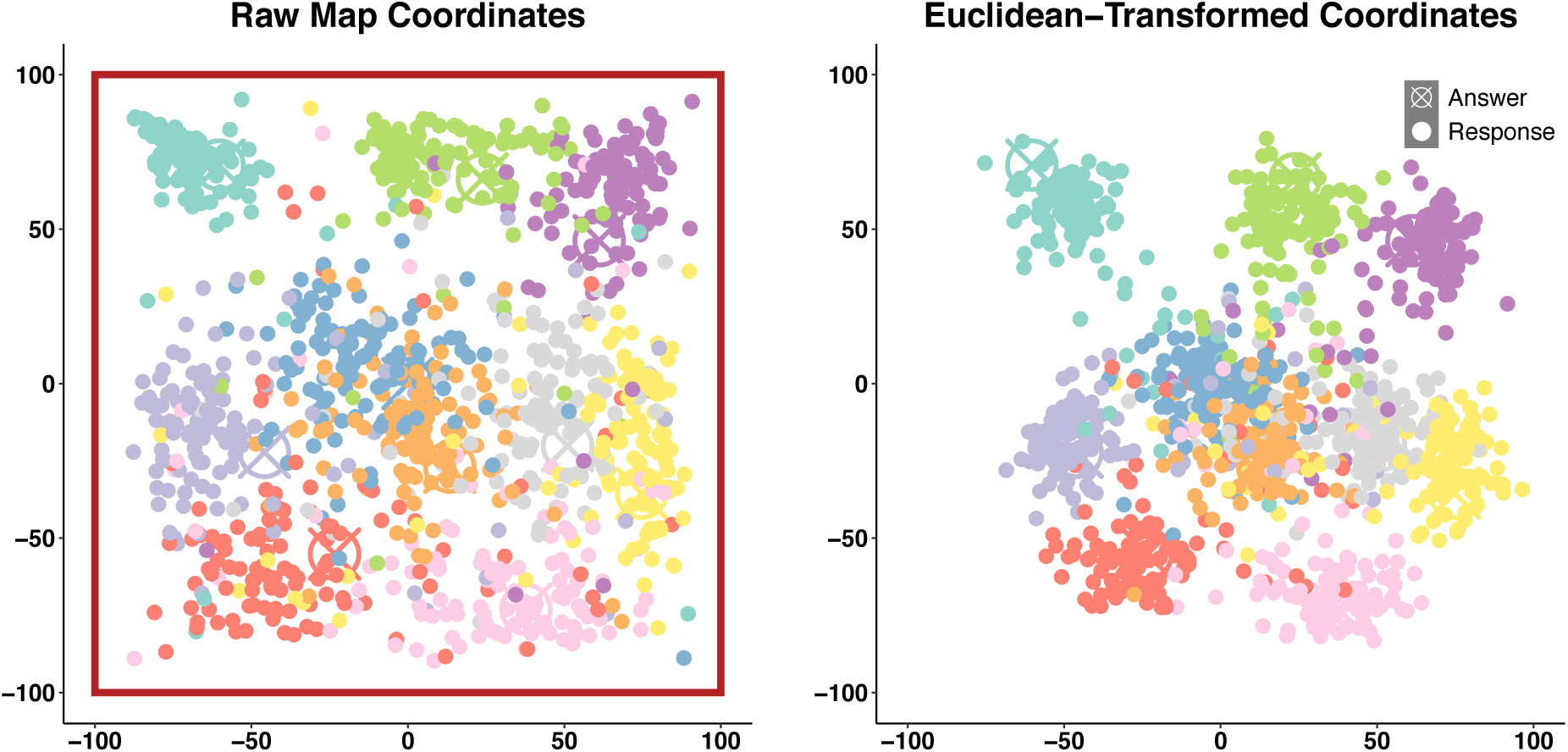
The participants’ maps were overall relatively accurate but displayed notable distortions. Left: The participants’ raw map responses (note: the red box depicts the locations of the boundaries of the environment; here, the participants drew maps that were significantly pulled toward the boundaries, terms a boundary expansion effect; also see Figure 18). Right: The Euclidean-transformed response coordinates. The circled x’s indicate the ground truth coordinates while the dots indicate individual participant coordinates and each location is depicted in a unique color.

**Figure 18:**
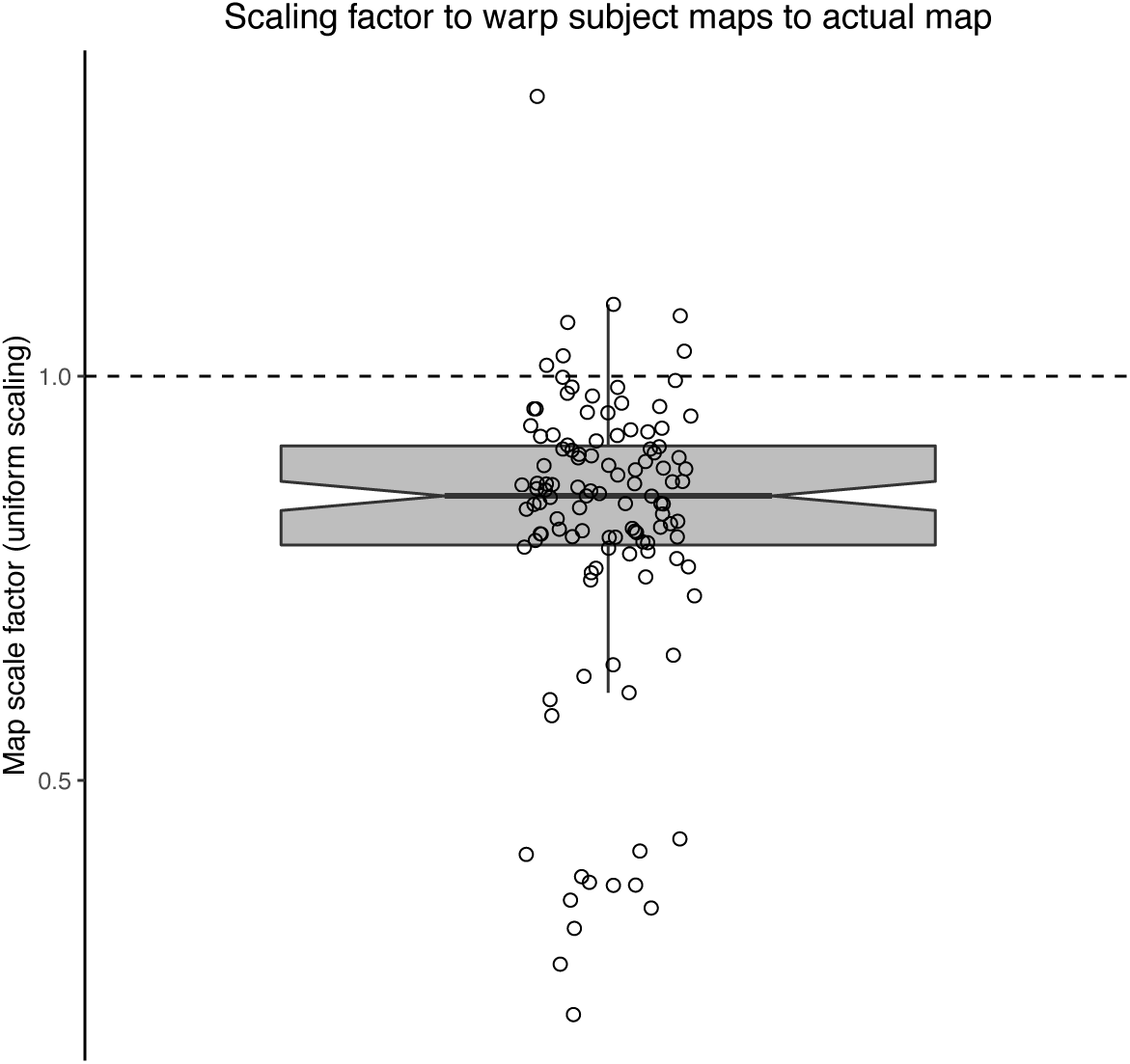
Evidence that participants warp their maps in a boundary-expansion effect (i.e., closer to the walls than the actual maps). Here, each dot indicates the (uniform) scaling factor required in the transformation from the participant map to the actual map. The boxplot depicts the middle 25^th^ to 75^th^ percentile of the data.

#### 6.2.2 Novel Virtual Environment: Evidence for Alignment Effects in Spatial Memory

As in the participants from Experiment 1, we next aimed to determine whether participants exhibited boundary alignment effects in their JRD responses for the virtual environment (see Figure 1B). Consistent with our predictions, we observed a significant effect of the alignment model parameter (alignment beta = 0.068, t = 3.99 p = 6.6 × 10^-5^; see Figure 19; note that the z-scored answer model parameter was also significant: answer beta = 0.18, t = 16.8, p < 2.0 × 10^-16^), thus replicating the boundary-alignment effect that we observed in Experiment 1 and in memory for the real-world environments.

**Figure 19:**
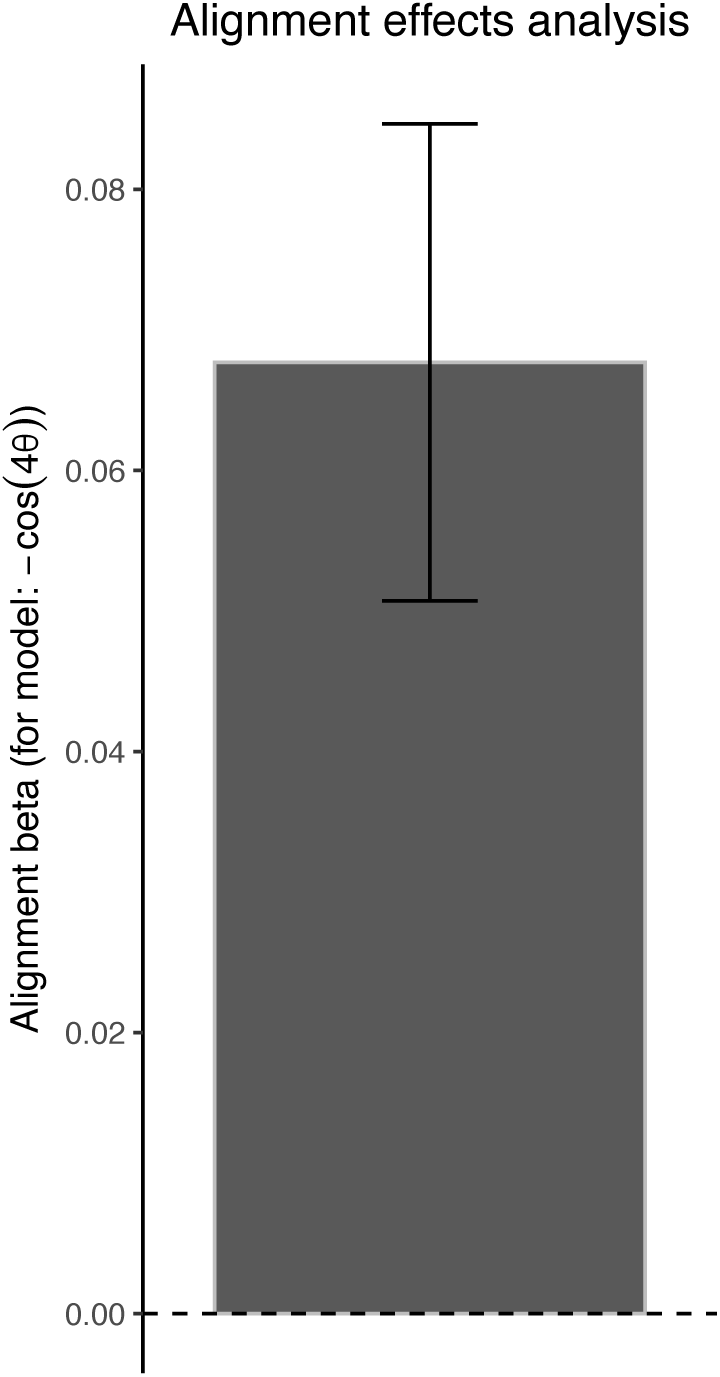
Evidence for an alignment effect within the JRD pointing accuracy (within the Experiment 2 data). The bar indicates the beta coefficient and the error bars indicate the standard errors from our mixed effects regression analysis.

#### 6.2.3 Virtual environment: Evidence that the JRD Task, the Map Task, and the Distance Estimation Task recruit common underlying representations

We again tested the relationship between the JRD Task, the Map Task, and the Distance Estimation Task with two analyses. First, we observed a significant relationship between performance on the JRD Task and the Map Task (Pearson’s *r* = −0.73, *t*_113_ = −11.34, *p* < 2.20 × 10^-16^, *BF*_10_ = 1.07 × 10^17^; Spearman’s *ρ* = −0.64, *S* = 416754, *p* < 2.20 × 10^-16^; Kendall’s *τ* = −0.45, *z* = −7.2, *p* = 5.99 × 10^-13^), the Distance Estimation Task and the Map Task (Pearson’s r = 0.67, t_110_ = 9.46, p = 6.72 × 10^−16^, BF_10_ = 6.70 × 10^12^; Spearman’s ρ = 0.66, S = 78776, p < 2.20 × 10^−16^; Kendall’s τ = 0.49, z = 7.65, p = 1.95 × 10^−14^; see Figure 19), and the JRD Task and the Distance Task (Pearson’s r = −0.7, t_110_ = −10.23, p < 2.20 × 10^−16^, BF_10_ = 3.05 × 10^14^; Spearman’s ρ = −0.7, S = 398268, p < 2.20 × 10^−16^; Kendall’s τ = 0.49, z = 7.65, p = 1.95 × 10^−14^; see Figure 20).

**Figure 20:**
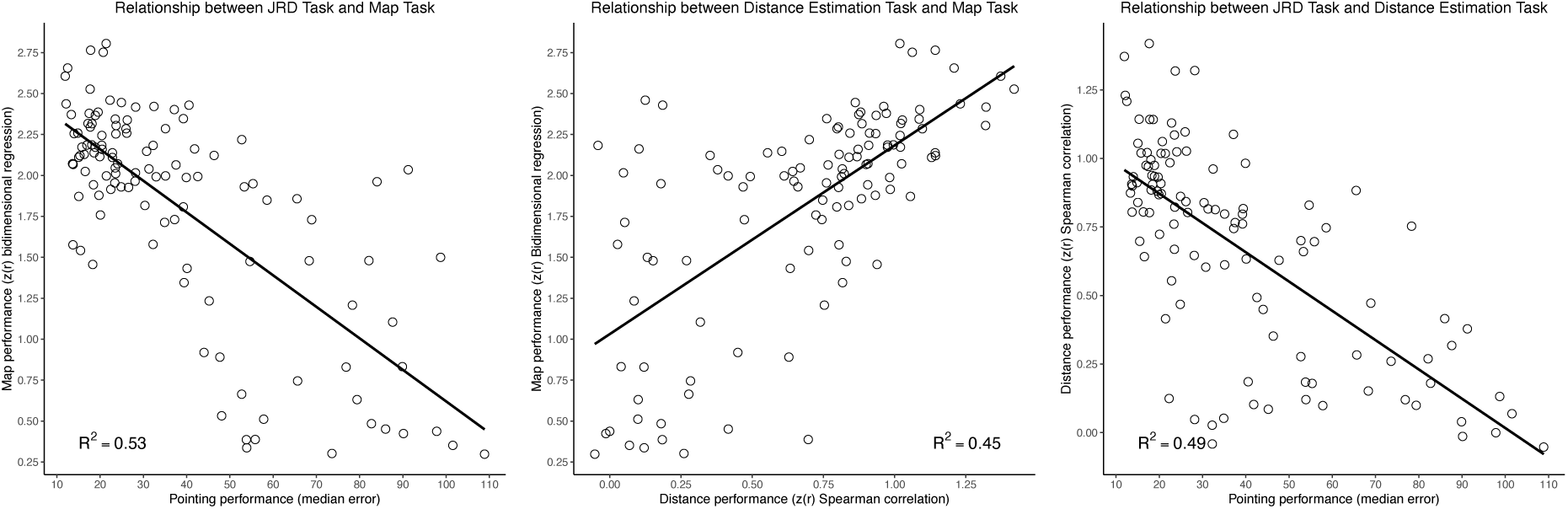
Relationship between the Map Task, JRD Task, and Distance Estimation Task for the virtual environment. Left: The relationship between performance on the JRD Task and the Map Task. Center: The relationship between performance on the Distance Estimation Task and the Map Task. Right: The relationship between performance on the JRD Task and the Distance Estimation Task.

Second, the patterns of errors were correlated across these tasks. Specifically, the Fisher’s r-to-z transformed partial circular correlation coefficients between angular responses on the JRD Task and the Map Task were significantly greater than zero when partialing out the angular answers (mean = 0.19, *t*_114_ = 8.55, *p* = 6.23 × 10^-14^, *BF*_10_ = 1.13 × 10^11^; see Figure 21). Likewise, the Fisher’s r-to-z transformed partial Spearman correlation coefficients between distances between landmarks on the Distance Estimation Task and the Map Task (while partialing out the effect of the answers) were significantly greater than zero mean = 0.31, *t*_111_ = 15.29, *p* < 2.20 × 10^-16^, *BF*_10_ = 7.73 × 10^25^; see Figure 21). The results of these correlation analyses provide further evidence that the JRD Task, the Map Task, and the Distance Estimation Task recruit similar underlying cognitive representations for the virtual environment, thus replicating the results from the real-world envirnments and the Experiment 1 data from the virtual environment (see above).

**Figure 21:**
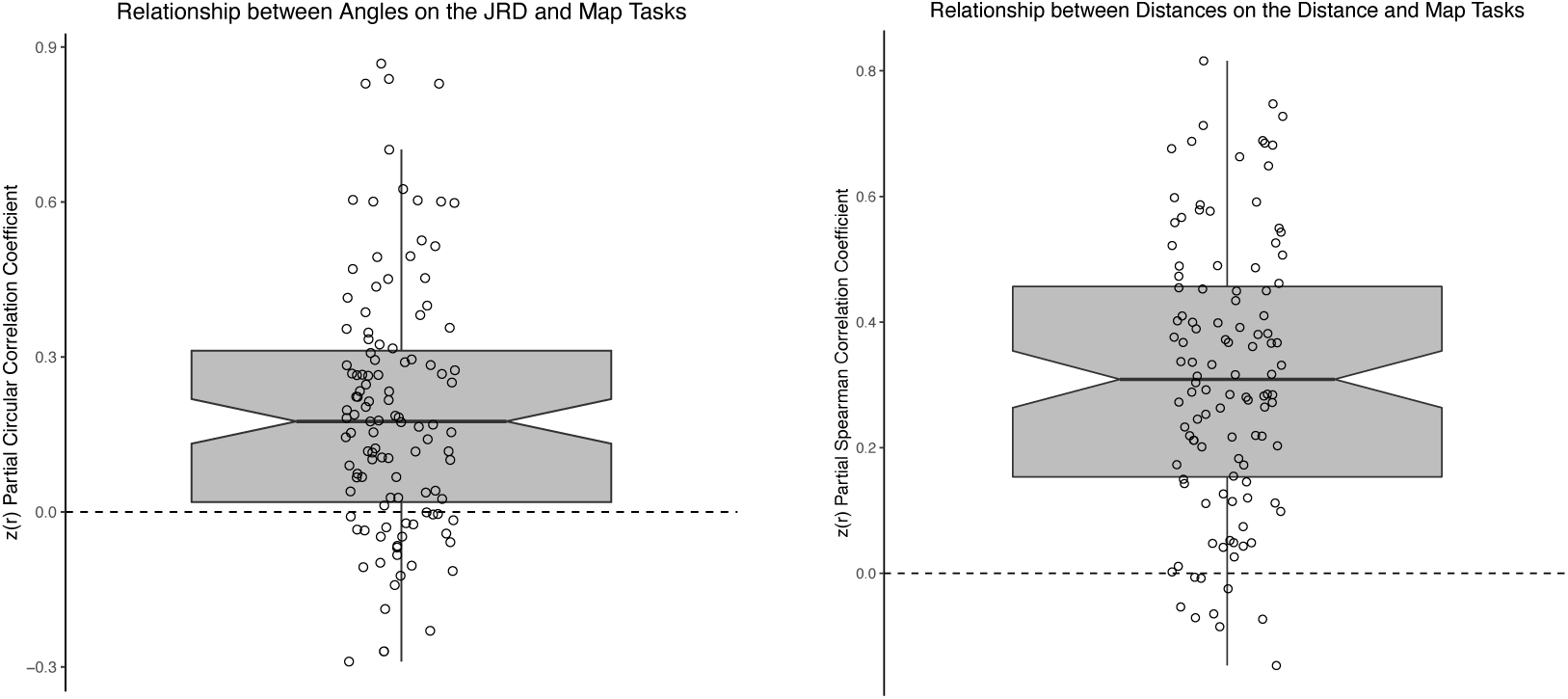
Evidence that participants’ recruit similar underlying cognitive representations to solve the JRD Task and the Map Task for the virtual environment. Left panel: The Fisher’s r-to-z transformed partial circular correlation coefficients for the responses to the Map Task and the JRD Task were significantly greater than zero (mean = 0.19, *t*_114_ = 8.56, *p* = 6.23 × 10^−14^, *BF*_10_ = 1.13 × 10^11^). Right panel: The Fisher’s r-to-z transformed partial Spearman correlation coefficients for the responses to the Map Task and the Distance Estimation Task were greater than zero (mean = 0.31, *t*_111_ = 15.29, *p <* 2.2 × 10^−16^, *BF*_10_ = 7.73 × 10^25^). Each dot indicates an individual participant and the boxplot depicts the middle 25th to 75th percentile of the data.

#### 6.2.4 Virtual environment: Evidence that learning is enhanced by repeated navigation

Our results reveal that pointing performance and confidence on the JRD Task improved with repeated rounds of navigation in the virtual environment (see Supplemental Information and Supplemental Figure 2), thus replicating the results of Experiment 1 and our previous work (Huffman & Ekstrom, 2019).

#### 6.2.5 Comparison between performance on the real-world (hometown) and virtual environment tasks

We next tested the correlations between performance on the real-world and virtual environment tasks. We found that the correlation between all three tasks was significant: JRD Task (Pearson’s *r* = 0.52, *t*_92_ = 5.82, *p* = 8.49 × 10^-8^, *BF*_10_ = 181126; Spearman’s *ρ* = 0.53, *S* = 64584, *p* = 5.19 × 10^-8^; Kendall’s *τ* = 0.36, *z* = 5.12, *p* = 2.10 × 10^-7^), Map Drawing Task (Pearson’s *r* = 0.26, *t*_92_ = 2.59, *p* = .011, *BF*_10_ = 4.93; Spearman’s *ρ* = 0.34, *S* = 91224, *p* = .00082; Kendall’s *τ* = 0.24, *z* = 3.38, *p* = .00072), and the Distance Estimation Task (Pearson’s *r* = 0.39, *t*_86_ = 3.91, *p* = .00018, *BF*_10_ = 178.1; Spearman’s *ρ* = 0.37, *S* = 71000, *p* = .00035; Kendall’s *τ* = 0.26, *z* = 3.63, *p* = .00028; see Figure 22). Altogether, these correlations show that there is certainly shared variance between performance on the real-world and virtual environments; however, as in Experiment 1, there is also a fair degree of unique variance (i.e., all of the r^2^’s < 0.4, thus explaining less than 40% of the variance), thus suggesting there may also be unique aspects of the real-world navigation task that cannot be fully captured with the novel virtual environment.

**Figure 22:**
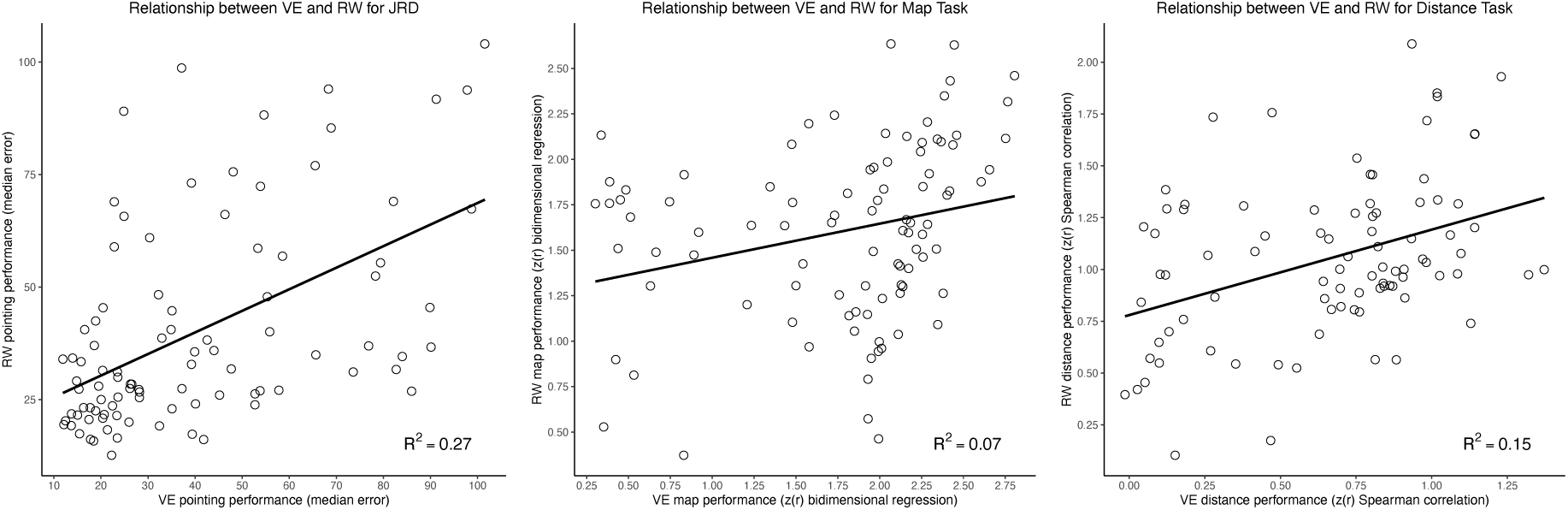
The correlations were significant for the comparison between performance on the virtual environment and the real world (hometown) for the JRD Task (see left panel), the Map Task (see middle panel), and the Distance Estimation Task (see right panel).

### 6.3 Discussion

Overall, the findings from the novel virtual environment task extend our previous findings, providing further evidence for non-Euclidean distortions. Specifically, we found that participants drew maps that were affected by a “boundary expansion effect.” The inclusion of the boundary in this experiment gave us a clear anchor point to determine the nature of memory distortions in map drawing. Moreover, we replicated the boundary alignment effect in the JRD data. We also observed correlations between performance on the tasks as well as the patterns of errors across tasks, thus suggesting partially overlapping spatial representations for JRD pointing, map drawing, and distance estimation. Moreover, we observed correlations between performance on the virtual and real-world environments. Altogether, our findings suggest that non-Euclidean distortions, due to heuristic use, are present across highly familiar, recently learned and ambulated, and novel large-scale environments.

## 7 General Discussion

As we discussed in the Introduction, there remains significant disagreement in the literature about the nature of the underlying spatial representations that we use to navigate and remember spatial environments. On the one hand, the Euclidean, cognitive map hypothesis has been highly influential in the neuroscientific literature. For example, the discovery of spatially tuned neurons, such as place cells and grid cells, led prominent theories to argue for a Euclidean representation of space (e.g., Bellmund et al., 2016; Gallistel, 1990; O’Keefe & Nadel, 1978).

On the other hand, based on decades of behavioral research in humans, other theories propose that the underlying representation of space is fuzzier and distorted, perhaps more akin to a cognitive graph, cognitive collage, or hierarchical representations of space, which cause systematic distortions and a lack of globally consistent metric knowledge (e.g., Chrastil & Warren, 2014; Du et al., 2023; Ekstrom et al., 2017; Ericson & Warren, 2020; Foo et al., 2005; McNamara, 1986, 1991; Tversky, 1981, 1992, 1993; Warren, 2019; Warren et al., 2017). However, the vast majority of support for the cognitive graph hypothesis comes from studies in which participants navigate and perform memory tasks for novel virtual environments, which may differ in terms of the complexity of the environment (e.g., in terms of the size, number of landmarks, layout of landmarks, duration of exposure to the environment, personal relevance of the environment). Thus, we aimed to determine whether we would observe similar effects across different environments, including large-scale, real-world environments that participants navigated over the course of months to many years. As we argue in the Introduction, if the systematic distortions extend to these environments and are consistently observed across multiple measures of memory, then we can at least partially rule out alternative explanations for systematic distortions in spatial memory (e.g., that participants did not have enough time or motivation to generate more accurate, globally consistent, Euclidean representations of space). Thus, we aimed to test between two highly influential theories of spatial memory. As we showed above and as we discuss in detail below, we provided evidence for systematic distortions in spatial memory across all of our environments and tasks.

Our map drawing results for the real-world environments provide one set of evidence for the hypothesis that remote memory for spatial environments is supported by systematic distorted representations that violate globally consistent Euclidean assumptions. For both the campus and hometown environments, we found strong evidence that an affine transformation better accounts for map-drawing performance than a so-called Euclidean transformation (see Figure 1A, Figure 2, and Figure 6). Because the affine transformation allowed for systematic distortions in their representations (i.e., shearing or scaling related to boundaries or landmarks, which can accommodate alignment and rotations heuristics; e.g., see Friedman & Kohler, 2003; Nakaya, 1997), our findings support the idea of systematic distortions in spatial representations. Importantly, our results build on previous research (Nakaya, 1997) by showing that the affine transformation better accounted for participants’ maps at various levels of exposure: from the college campus, which is explored over the period of months to years, to the hometown environment, which is explored for years to decades. Furthermore, while Nakaya (1997) averaged the coordinates across participants before performing the Bidimensional Regression analysis, we instead ran the analysis separately within each participant and then compared the aggregate ΔAIC values at the group level. Thus, our approach is more akin to a random effects analysis vs. a fixed effects analysis and helps rule out the possibility that the previous results were driven by an artifact of averaging the map locations across participants prior to the analysis. Moreover, the findings in the hometown environment are important because they show that these effects generalize across multiple environments and are not restricted to any specific environment or subset of participants. Altogether, our findings support the idea that human spatial representations violate Euclidean assumptions, even when they are well-learned and highly familiar.

The results of the map drawing analyses for the novel virtual environment provide evidence for systematic distortions in spatial representations of novel environments, thus suggesting that some of the same distortions are present as for familiar, well learned environments. Importantly, the boundary of the environment was visible to the participants during the map-drawing task, which allowed us to further interrogate the specific nature of distortions in participants’ maps. Here, we found evidence that participants exhibited a “boundary expansion effect,” in which they drew the outer landmarks as closer to the boundaries than they actually were (see Figure 1D, Figure 11, Figure 12, Figure 17, Figure 18). These findings suggest that participants employed a key heuristic such that they anchored the representations to the boundaries of the environment. Moreover, we found evidence that the “boundary expansion effect” led participants to also draw the outer landmarks as more aligned to the boundaries than they actually were (see Supplemental Figure 3). Altogether, these results provide additional information about the nature of spatial memory distortions and provide novel insight into memory heuristics, thus building on previous studies that have highlighted a key role for environmental boundaries on spatial memory (Brunec, Moscovitch, et al., 2018; Doeller & Burgess, 2008; Miller et al., 2018). We think it will be interesting for future studies to include similar boundary-like information in real-world environments to determine whether similar effects are seen for more remote spatial memory, including real-world environments.

The alignment effects in the JRD task (see Figure 1B) provide additional evidence for systematic distortions in spatial memory for both novel and remote spatial environments. Specifically, we found significant evidence of boundary alignment effects in JRD pointing accuracy for the campus environment (see Figure 3), the hometown environment (see Figure 8), and the novel virtual environment (see Figure 13 and Figure 19). Our findings in the novel virtual environment replicate and extend upon on previous research in novel environments. For example, previous research has found alignment effects for new learning within small-scale, real-world environments (Mou et al., 2006, 2007; Shelton & McNamara, 1997, 2001), large-scale, real-world environments (McNamara et al., 2003), and large-scale virtual environments (Starrett et al., 2019). Thus, there is a confluence of evidence across paradigms to support the idea that novel spatial learning is subjected to boundary alignment effects in JRD pointing. However, while (Marchette et al., 2011) found evidence of boundary alignment effects for participants’ university campus, much less was hitherto known about whether boundary alignment effects would persist for very well learned, personally relevant, real-world spatial environments or how these representations may change with more time for learning. Thus our finding of significant alignment effects for the campus task replicates the previous findings (Marchette et al., 2011). Moreover, the extension to the hometown environment suggests that these effects persist for environments that are explored for an even longer period of time and with even more individual variability in terms of the actual physical environment itself (i.e., since participants had largely unique hometown environments), thus ruling out possible alternative explanations about whether the findings are particular for a given chosen environment. Altogether, our findings suggest that the boundary alignment effect in JRD pointing responses is a relatively ubiquitous property of human spatial memory. It will be interesting for future experiments to replicate these findings and to determine whether there are any conditions in which alignment effects are not present, but, at the present time, we conclude that spatial memory distortions appear to be prevalent across both novel and remote scales.

Importantly, we found that both individual performance and the pattern of errors across the map-drawing task, JRD task, and distance estimation task were all correlated, thus suggesting that all of the tasks tap into partially overlapping representations. These results are important because they suggest that the systematic distortions that we observed are not driven solely by nuances of any specific task, thus supporting the notion of converging operations (Garner et al., 1956; McNamara, 1991). Additionally, the significant correlation between patterns of errors argues strongly against the notion that participants might use multiple representations for each task, thus allowing us to sidestep concerns regarding generalization between tasks. Specifically, across multiple tasks, the representations share significant variance of the errors, thus suggesting that the similar underlying systematically distorted representations is used for all tasks. Our findings replicate and extend upon our previous research (Huffman & Ekstrom, 2019), and further support the idea that coarse knowledge of direction and distance are stored within a shared spatial representation, which is consistent across both novel and remote memory for both virtual and real-world environments. In summary, our findings support the notion that systematic distortions are a ubiquitous property of human spatial memory, even for very well learned and personally relevant environments.

Overall, our results support the cognitive graph hypothesis and provide a novel challenge to the Euclidean, cognitive map hypothesis. The finding of grid cell and grid-cell-like coding has led some prominent theories to suggest that the apparent metric nature of the neural responses in the medial entorhinal cortex would manifest as metric behavior for navigation (Hafting et al., 2005; Moser & Moser, 2008) and some have further posited grid-cell-like coding would play a role non-navigation-related cognition more generally (e.g., Behrens et al., 2018; Bellmund et al., 2018; Constantinescu et al., 2016; Park et al., 2021). However, we would like to emphasize that these theories have often been advanced on the basis of patterns of neural signals rather than on the basis of behavioral performance and recent theories have questioned how well the neural findings would map onto behavior, especially for large-scale spatial environments (e.g., Ekstrom et al., 2017, 2020; Ginosar et al., 2023; Stella et al., 2020).

We note that the cognitive graph hypothesis can account for a range of findings in the literature, including findings that were initially taken to support the cognitive map hypothesis. For example, shortcuts could be driven by landmark and familiarity-based navigation or forms of egocentric navigation (Bennett, 1996; Cruse & Wehner, 2011; Ekstrom et al., 2014). Previous computational modeling work also provides important support for non-globally consistent and more graph-based navigation strategies. For example, Cruse & Wehner (2011) created an artificial neural network that was able to perform novel shortcuts based on path integration alone (i.e., knowledge of the general direction of the goal, allowing you to figure out a more direct path), simulating the navigation behavior of ants which was previously a strong foundation for the use of cognitive map in animals and insects. Moreover, other recent computational modeling work with the successor representation demonstrated that it can account for distortions that occur to both grid-cell firing in rodents and behavioral error patterns in humans (Bellmund et al., 2019). A very recent theoretical paper argued that the successor representation can be employed as a tool to generate testable predictions of the cognitive graph hypothesis, specifically proposing that primate navigation is heavily influenced by visual information (Huffman et al., 2025). Additionally, Huffman et al. (2025) demonstrated that the model can account for several findings both from the neuroscientific and behavioral literatures, thus building on previous influential work with the successor representation (e.g., Bellmund et al., 2019; Stachenfeld et al., 2017). Given Huffman et al.’s core hypothesis of vision as playing a predominant role in primate spatial navigation (also see Ekstrom, 2015; Martinez-Trujillo, 2025; Martinez-Trujillo & Piza, 2026; Rolls, 2023), we can further interpret our findings here as suggesting that “just good enough” representations (cf. Ekstrom et al., 2017, 2020) from abstract spatial memory (e.g., pointing to unseen targets or drawing a map from memory) can perhaps serve as rough templates for more accurate wayfinding when people are situated within the environment (e.g., when they can employ additional strategies such as piloting or beaconing around familiar landmarks or key decision points). Thus, we think that it will be interesting for future research to generate further predictions from the cognitive graph hypothesis within the modeling framework to further expand on our results here.

In conclusion, we provide novel evidence that spatial memory for both novel and remote environments is supported by an underlying cognitive graph. We provide evidence for systematic distortions in both map drawing (e.g., affine transformation, “boundary expansion effect”, boundary alignment effect) and JRD pointing (e.g., boundary alignment effects) as well as correlations between patterns of errors between tasks. Altogether, our converging results suggest that systematic distortions are a ubiquitous property of human spatial memory across recent and remote timescales, including very-well learned environments that have daily significance and relevance to participants, thus raising an important challenge for models that propose a globally consistent, Euclidean-based view of spatial memory.

## 9 Appendix

### 9.1 Simulations of the partial circular correlation technique for analyzing the similarity of the errors on the JRD Task and the Map Task

We ran several simulations to substantiate the validity of the partial circular correlation technique. We were interested in determining the specificity (e.g., verifying that there were not unexpected effects driving our results due to assumptions of circular correlation) and sensitivity (e.g., that significant effects would be observed when there are significant correlations in the pattern of errors) of the partial correlation approach.

#### 9.1.1 We do not observe significant partial correlations when we simulate responses as the answers plus randomly distributed noise

We first wanted to rule out the possibility that our partial circular correlation results could be explained by a response model in which participants select their responses on the JRD Task based on the answer plus randomly distributed noise. Specifically, one idea could be that the magnitude or directions of the angles could somehow artificially induce significant partial circular correlations between the JRD Task and the Map Task. Thus, we simulated responses as the answers plus random noise by adding values from a von Mises distribution. Importantly, our simulations revealed that these simulations did not result in significant partial circular correlation results.

#### 9.1.2 We observed a significant partial circular correlation coefficient when the simulated patterns of errors were correlated

Importantly, we validated that our approach would reveal significant effects when the patterns of error were correlated between the two tasks. Specifically, for each trial, we generated correlated errors by creating two noise sources using von Mises distributions (noise distribution 1: kappa density parameter = X; noise distribution 2: kappa density parameter = X + Y; i.e., the density parameter is larger for the second noise distribution and is thus results in a tighter distribution of the data). We then simulated responses on the two tasks by adding the answer distribution to the first noise distribution (for Task 1) and by adding the answer distribution plus the two noise distributions (for Task 2). Note, because the spread of the distribution of the second noise distribution is smaller than the first noise distribution, this results in correlated “errors.” We observed a significant partial circular correlation coefficient between these two simulated responses. These results suggest that our approach appropriately detects a partial circular correlation coefficient under conditions with correlated errors, thus bolstering our interpretation of our empirical data.

## Author Contributions

Derek J. Huffman played a sole role in data curation, formal analysis, funding acquisition, investigation, methodology, project administration, resources, software, supervision, and visualization as well as a lead role in conceptualization, writing–original draft, and writing–review and editing. Arne D. Ekstrom played a supporting role in conceptualization, writing–original draft, and writing–review and editing. Nikhil Jaha played a lead role in data collection and a supporting role in conceptualization.

## Generative AI Disclosure Statement

We did not use any generative AI (e.g., large language models) in writing this paper (i.e., we wrote the entire manuscript on our own).

## Acknowledgements

Derek J. Huffman was partially supported by a grant from the National Institute of Mental Health of the National Institutes of Health (grant number: F32MH116577; during data collection and preliminary data analysis) as well as startup and other funds from the Department of Psychology and Colby College (continued and final data analysis, writing, etc.). Arne D. Ekstrom was supported by grants from the National Science Foundation (Behavioral and Cognitive Sciences Division grant number: NSF BCS-1630296) and the National Institute of Neurological Disorders and Stroke of the National Institutes of Health (grant number: R01NS076856). For the supplemental analysis of the warped angles in the Map Drawing Task from the virtual environment (see Supplemental Material and Supplemental Figure 3), we thank Chandrachud Gowda for implementing the analysis (under the guidance of DJH) and Jolien Cremers, the developer of the bpnreg package in R, for providing instructions for using the *bpnme* function. Finally, we also thank members of the Huffman Environmental Cognition Lab (especially Ainsley Bonin), the Human Spatial Cognition Lab (especially Michael Starrett), and the Dynamic Memory Lab (especially Charan Ranganath, Trevor Baer, and Angelique Delarazan) for helpful conversations and for providing logistical support for these experiments.

## Supplemental Material

Here, we report additional methods, results, and figures from the data from both experiments for the virtual environment, including the JRD pointing performance and confidence over blocks of learning/testing (here participants completed 6 blocks that alternated between the navigation task and the JRD Task; see section S1 and Supplemental Figures 1 and 2) as well as an analysis of boundary alignment effects in the map drawing data (see section S2 and Supplemental Figure 3).

### S1 Additional Analysis of the Evolution of Performance on the JRD Task over Blocks of Learning in the Novel Virtual Environment

Our results reveal that pointing performance on the JRD Task improved with repeated rounds of navigation in the virtual environment in both Experiment 1 (see Supplemental Figure 1) and Experiment 2 (see Supplemental Figure 2). Interestingly, however, there was a large spread in performance across blocks, where some participants reached peak performance after a single block (e.g., median absolute angular error less than 20 degrees) while other participants started with a high degree of error and never improved across blocks (note that there were 20 trials of the JRD Task for the first 5 blocks and 60 for the final block). These results are consistent with previous studies that have found a large degree of individual differences in spatial memory performance over the course of learning (e.g., Ishikawa & Montello, 2006; please see the main text for further discussion). Relatedly, memory strength (as assessed by confidence) increased over blocks of the task; however, there were several participants that maintained low levels of confidence throughout all blocks of the task. Our findings here also replicate our previous work (Huffman & Ekstrom, 2019).

**Supplemental Figure 1:**
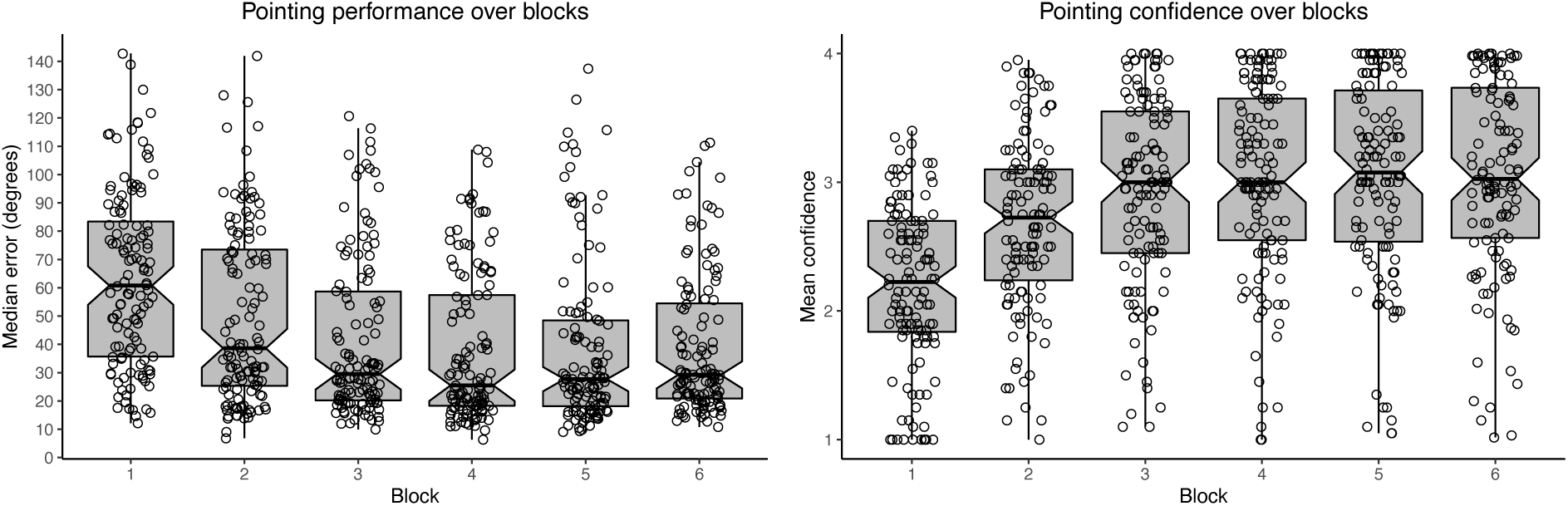
Evidence that repeated navigation in the virtual environment improves JRD pointing performance (left panel) and increases confidence (right panel; data from Experiment 1).

**Supplemental Figure 2:**
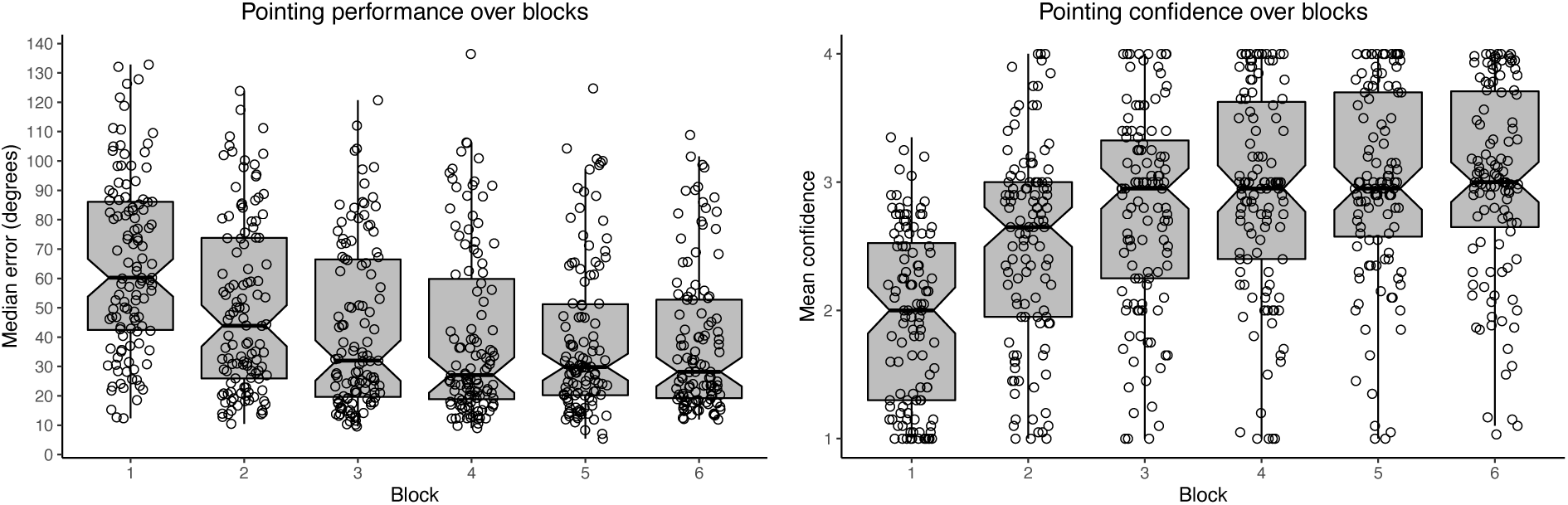
Evidence that repeated navigation in the virtual environment improves JRD pointing performance (left panel) and increases confidence (right panel; data from Experiment 2).

### S2 Additional Analysis of the Warping of Angles in the Map Drawing Task

We were next interested in following up on the “boundary expansion effect,” in which participants drew outer landmarks as closer to the boundaries than they actually were (see main text and Figure 11, Figure 12, Figure 17, and Figure 18) by testing whether participants tended to draw their maps with the outer landmarks relatively more aligned than they actually were. Specifically, the cognitive graph hypothesis would predict that errors would be biased, such that they are more aligned with the main axes of the environment than they were in reality. In contrast, the Euclidean cognitive map hypothesis would predict that errors would be randomly distributed around the true answer (i.e., there would be error, but that error should be randomly distributed and not systematically biased, thus being closer to a Gaussian distribution around the true answer). Thus, to test between these two predictions, we implemented a circular analysis using the function *bpnme* from the package bpnreg (Cremers & Klugkist, 2018) within R to determine if the angles of the outer stores were more aligned than they were in the actual environment. Here, we transformed the data such that participant angles that were pulled in the direction of the aligned angle were always positive (and those that were pulled in the direction away from the aligned angle were negative), thus any positive skew would indicate that participants tended to draw the outer landmarks as more aligned than they actually were (consistent with the cognitive graph hypothesis), whereas a null angle that included 0 in the confidence interval would indicate that their angles were relatively evenly distributed about 0 and were not significantly biased by the alignment geometry of the environment (consistent with the globally consistent, Euclidean, cognitive map hypothesis). Given that we were only interested in determining if the mean angles were pulled toward a significant map/boundary alignment effect (i.e., we were not testing for any main effects or interactions), we implemented an intercepts model:

~~~
fit_intercept <- bpnme(
  pred.I = Phaserad ∼ 1 + (1 | ParticipantNumber),
  data = angle_data,
  its = 10000,
  burn = 1000,
  n.lag = 3,
  seed = 101
)
~~~

**Supplemental Figure 3:**
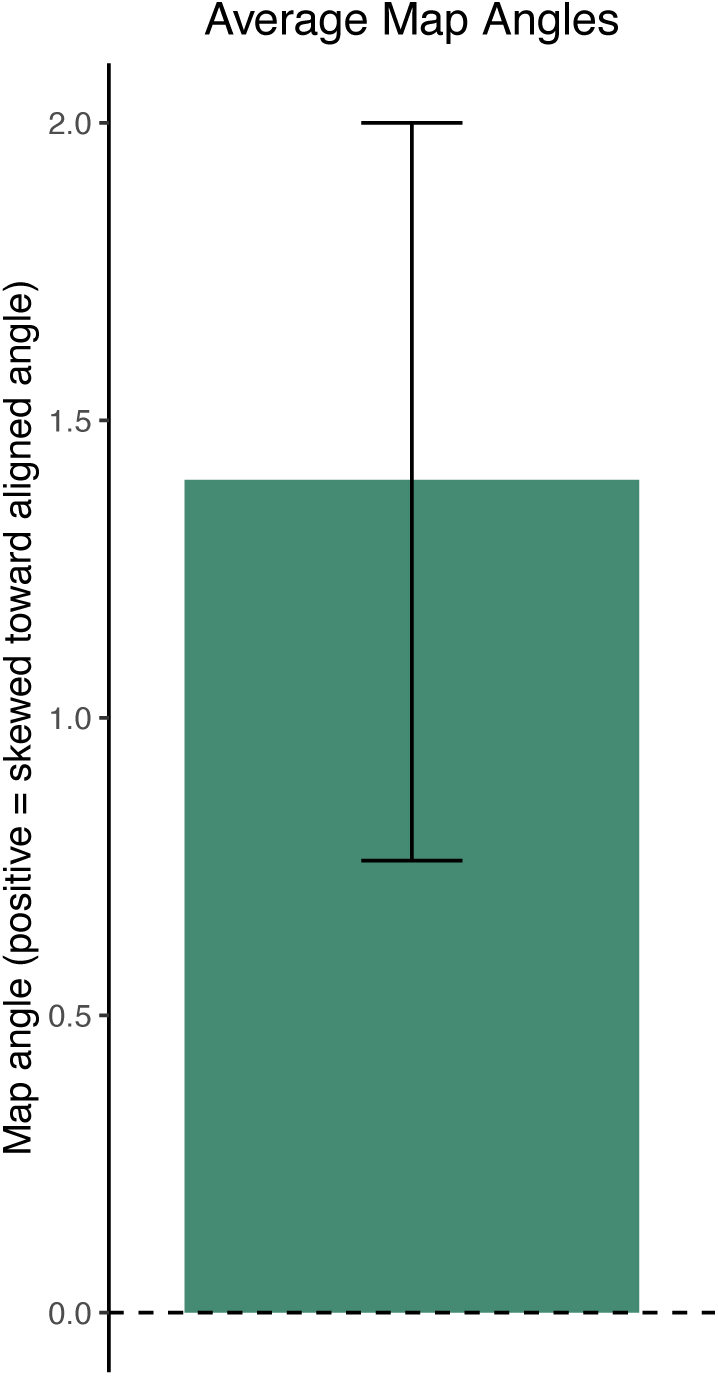
Evidence that participant’s warp their map data for the outer landmarks to be more aligned along the outer walls than they were in reality (data from the virtual environments in Experiment 1 and 2). The bar indicates the mean and the error bars are the 95% HPD confidence intervals (from the output of *bpnme*). We found that the errors were significantly positive, indicating that participants tended to skew their map drawing performance to be more aligned than the actual answers for the outer landmarks in the virtual environment task.

Note that *bpnme* is a Bayesian approach and we can gather the 95% highest posterior density interval (HPD). As Cremers and Klugkist (2018) describe, the HPD intervals allow statements about probability that an effect lies within a given range and thus serve as a test for significance (in our case, the estimate for the intercept term). Given that there were no theoretical differences between the data from the virtual environments in both experiments, we combined the data here to have maximal power to detect an effect. Consistent with our prediction of a map alignment effect, we found that the angle at which the participants drew the outer landmarks was significantly skewed toward the aligned angle (i.e., rather than evenly distributed around the true answer, which would be predicted by the Euclidean cognitive maps hypothesis): mean = 1.4 degrees; HPD interval = 0.8 to 2.0 (note that the HPD interval is a measure for statistical significance at the 95% level, since it indicates the 95% probability range for a given observed effect; see our results in Supplemental Figure 3; for more information about this statistical approach, please see Cremers & Klugkist, 2018).

Altogether, our results here build on the “boundary expansion effect” by suggesting that as participants draw outer landmarks as closer to the environmental boundaries, they tend to also draw the outer landmarks as more aligned than they actually were. Therefore, our results provide important additional insight into the nature of systematic distortions in spatial memory judgments, further supporting the cognitive graph hypothesis.

## References

Behrens, T. E. J., Muller, T. H., Whittington, J. C. R., Mark, S., Baram, A. B., Stachenfeld, K. L., & Kurth-Nelson, Z. (2018). What Is a Cognitive Map? Organizing Knowledge for Flexible Behavior. Neuron, 100(2), 490–509. 10.1016/j.neuron.2018.10.002

Bellmund, J. L., Deuker, L., Navarro Schröder, T., & Doeller, C. F. (2016). Grid-cell representations in mental simulation. eLife, 5, e17089. 10.7554/eLife.17089

Bellmund, J. L. S., de Cothi, W., Ruiter, T. A., Nau, M., Barry, C., & Doeller, C. F. (2019). Deforming the metric of cognitive maps distorts memory. Nature Human Behaviour, 4(2), 177–188. 10.1038/s41562-019-0767-3

Bellmund, J. L. S., Gärdenfors, P., Moser, E. I., & Doeller, C. F. (2018). Navigating cognition: Spatial codes for human thinking. Science, 362(6415), eaat6766. 10.1126/science.aat6766

Bennett, A. T. (1996). Do animals have cognitive maps? The Journal of Experimental Biology, 199(Pt 1), 219–224.

Boeing, G. (2017). OSMnx: New methods for acquiring, constructing, analyzing, and visualizing complex street networks. Computers, Environment and Urban Systems, 65, 126–139. 10.1016/j.compenvurbsys.2017.05.004

Bonin, A. K., & Huffman, D. J. (2026). Map or Graph? Systematic Biases Inform the Structure of Human Spatial Representations. PsyArXiv.

Brunec, I. K., Bellana, B., Ozubko, J. D., Man, V., Robin, J., Liu, Z.-X., Grady, C., Rosenbaum, R. S., Winocur, G., Barense, M. D., & Moscovitch, M. (2018). Multiple Scales of Representation along the Hippocampal Anteroposterior Axis in Humans. Current Biology, 28(13), 2129–2135.e6. 10.1016/j.cub.2018.05.016

Brunec, I. K., Moscovitch, M., & Barense, M. D. (2018). Boundaries Shape Cognitive Representations of Spaces and Events. Trends in Cognitive Sciences, 22(7), 637–650. 10.1016/j.tics.2018.03.013

Carbon, C.-C. (2013). BiDimRegression: Bidimensional Regression Modeling Using *R*. Journal of Statistical Software, 52(Code Snippet 1). 10.18637/jss.v052.c01

Chrastil, E. R., & Warren, W. H. (2013). Active and passive spatial learning in human navigation: Acquisition of survey knowledge. Journal of Experimental Psychology. Learning, Memory, and Cognition, 39(5), 1520–1537. 10.1037/a0032382

Chrastil, E. R., & Warren, W. H. (2014). From cognitive maps to cognitive graphs. PloS One, 9(11), e112544. 10.1371/journal.pone.0112544

Clemenson, G. D., Henningfield, C. M., & Stark, C. E. L. (2019). Improving Hippocampal Memory Through the Experience of a Rich Minecraft Environment. Frontiers in Behavioral Neuroscience, 13, 57. 10.3389/fnbeh.2019.00057

Clemenson, G. D., & Stark, C. E. L. (2015). Virtual Environmental Enrichment through Video Games Improves Hippocampal-Associated Memory. Journal of Neuroscience, 35(49), 16116–16125. 10.1523/JNEUROSCI.2580-15.2015

Constantinescu, A. O., O’Reilly, J. X., & Behrens, T. E. J. (2016). Organizing conceptual knowledge in humans with a gridlike code. Science, 352(6292), 1464–1468. 10.1126/science.aaf0941

Coutrot, A., Manley, E., Goodroe, S., Gahnstrom, C., Filomena, G., Yesiltepe, D., Dalton, R. C., Wiener, J. M., Hölscher, C., Hornberger, M., & Spiers, H. J. (2022). Entropy of city street networks linked to future spatial navigation ability. Nature, 604(7904), 104–110. 10.1038/s41586-022-04486-7

Cremers, J., & Klugkist, I. (2018). One Direction? A Tutorial for Circular Data Analysis Using R With Examples in Cognitive Psychology. Frontiers in Psychology, 9, 2040. 10.3389/fpsyg.2018.02040

Cruse, H., & Wehner, R. (2011). No Need for a Cognitive Map: Decentralized Memory for Insect Navigation. PLoS Computational Biology, 7(3), e1002009. 10.1371/journal.pcbi.1002009

Doeller, C. F., & Burgess, N. (2008). Distinct error-correcting and incidental learning of location relative to landmarks and boundaries. Proceedings of the National Academy of Sciences, 105(15), 5909–5914. 10.1073/pnas.0711433105

Du, Y. K., McAvan, A. S., Zheng, J., & Ekstrom, A. D. (2023). Spatial memory distortions for the shapes of walked paths occur in violation of physically experienced geometry. PLOS ONE, 18(2), e0281739. 10.1371/journal.pone.0281739

Ekstrom, A. D. (2015). Why vision is important to how we navigate: Human Spatial Navigation and Vision. Hippocampus, 25(6), 731–735. 10.1002/hipo.22449

Ekstrom, A. D., Arnold, A. E. G. F., & Iaria, G. (2014). A critical review of the allocentric spatial representation and its neural underpinnings: Toward a network-based perspective. Frontiers in Human Neuroscience, 8. 10.3389/fnhum.2014.00803

Ekstrom, A. D., Harootonian, S. K., & Huffman, D. J. (2020). Grid coding, spatial representation, and navigation: Should we assume an isomorphism? Hippocampus, 30(4), 422–432. 10.1002/hipo.23175

Ekstrom, A. D., Huffman, D. J., & Starrett, M. (2017). Interacting networks of brain regions underlie human spatial navigation: A review and novel synthesis of the literature. Journal of Neurophysiology, 118(6), 3328–3344. 10.1152/jn.00531.2017

Ekstrom, A. D., & Isham, E. A. (2017). Human spatial navigation: Representations across dimensions and scales. Current Opinion in Behavioral Sciences, 17, 84–89. 10.1016/j.cobeha.2017.06.005

Ekstrom, A. D., Spiers, H. J., Bohbot, V. D., & Rosenbaum, R. S. (2018). Human spatial navigation. Princeton University Press.

Epstein, R. A., Patai, E. Z., Julian, J. B., & Spiers, H. J. (2017). The cognitive map in humans: Spatial navigation and beyond. Nature Neuroscience, 20(11), 1504–1513. 10.1038/nn.4656

Ericson, J. D., & Warren, W. H. (2020). Probing the invariant structure of spatial knowledge: Support for the cognitive graph hypothesis. Cognition, 200, 104276. 10.1016/j.cognition.2020.104276

Foo, P., Warren, W. H., Duchon, A., & Tarr, M. J. (2005). Do Humans Integrate Routes Into a Cognitive Map? Map- Versus Landmark-Based Navigation of Novel Shortcuts. Journal of Experimental Psychology: Learning, Memory, and Cognition, 31(2), 195–215. 10.1037/0278-7393.31.2.195

Frankenstein, J., Mohler, B. J., Bülthoff, H. H., & Meilinger, T. (2012). Is the Map in Our Head Oriented North? Psychological Science, 23(2), 120–125. 10.1177/0956797611429467

Friedman, A., & Kohler, B. (2003). Bidimensional Regression: Assessing the Configural Similarity and Accuracy of Cognitive Maps and Other Two-Dimensional Data Sets. Psychological Methods, 8(4), 468–491. 10.1037/1082-989X.8.4.468

Gagnon, S. A., Brunyé, T. T., Gardony, A., Noordzij, M. L., Mahoney, C. R., & Taylor, H. A. (2014). Stepping Into a Map: Initial Heading Direction Influences Spatial Memory Flexibility. Cognitive Science, 38(2), 275–302. 10.1111/cogs.12055

Gallistel, C. R. (1990). The organization of learning. the MIT press.

Garner, W. R., Hake, H. W., & Eriksen, C. W. (1956). Operationism and the concept of perception. Psychological Review, 63(3), 149–159. 10.1037/h0042992

Ginosar, G., Aljadeff, J., Las, L., Derdikman, D., & Ulanovsky, N. (2023). Are grid cells used for navigation? On local metrics, subjective spaces, and black holes. Neuron, 111(12), 1858–1875. 10.1016/j.neuron.2023.03.027

Hafting, T., Fyhn, M., Molden, S., Moser, M.-B., & Moser, E. I. (2005). Microstructure of a spatial map in the entorhinal cortex. Nature, 436(7052), 801–806. 10.1038/nature03721

Hirtle, S. C., & Jonides, J. (1985). Evidence of hierarchies in cognitive maps. Memory & Cognition, 13(3), 208–217. 10.3758/BF03197683

Huffman, D. J., Dang, L. K., Doherty, O. R., Shakya, A., & Bonin, A. (2025). All Eyes on the Hippocampus: The Primate Hippocampus as a Visually-Guided Cognitive Graph. Proceedings of the Annual Meeting of the Cognitive Science Society, 47. https://escholarship.org/uc/item/8444g7k8

Huffman, D. J., & Ekstrom, A. D. (2019). Which way is the bookstore? A closer look at the judgments of relative directions task. Spatial Cognition & Computation, 19(2), 93–129. 10.1080/13875868.2018.1531869

Ishikawa, T., & Montello, D. (2006). Spatial knowledge acquisition from direct experience in the environment: Individual differences in the development of metric knowledge and the integration of separately learned places. Cognitive Psychology, 52(2), 93–129. 10.1016/j.cogpsych.2005.08.003

Julian, J. B., Keinath, A. T., Frazzetta, G., & Epstein, R. A. (2018). Human entorhinal cortex represents visual space using a boundary-anchored grid. Nature Neuroscience, 21(2), 191–194. 10.1038/s41593-017-0049-1

Krzanowski, W. J. (2000). Principles of Multivariate Analysis: A User’s Perspective. Oxford University PressOxford. 10.1093/oso/9780198507086.001.0001

Marchette, S. A., Yerramsetti, A., Burns, T. J., & Shelton, A. L. (2011). Spatial memory in the real world: Long-term representations of everyday environments. Memory & Cognition, 39(8), 1401–1408. 10.3758/s13421-011-0108-x

Martinez-Trujillo, J. (2025). Why do primates have view cells instead of place cells? Trends in Cognitive Sciences, 29(3), 226–229. 10.1016/j.tics.2024.12.007

Martinez-Trujillo, J., & Piza, D. (2026). The Visual Umwelt of primates and Hippocampal Representations of Space. Hippocampus, 36(1), e70053. 10.1002/hipo.70053

McNamara, T. P. (1986). Mental representations of spatial relations. Cognitive Psychology, 18(1), 87–121. 10.1016/0010-0285(86)90016-2

McNamara, T. P. (1991). Memory’s View of Space. In Psychology of Learning and Motivation (Vol. 27, pp. 147–186). Elsevier. 10.1016/S0079-7421(08)60123-1

Mcnamara, T. P., & Diwadkar, V. A. (1997). Symmetry and Asymmetry of Human Spatial Memory. Cognitive Psychology, 34(2), 160–190. 10.1006/cogp.1997.0669

McNamara, T. P., Hardy, J. K., & Hirtle, S. C. (1989). Subjective hierarchies in spatial memory. Journal of Experimental Psychology: Learning, Memory, and Cognition, 15(2), 211–227. 10.1037/0278-7393.15.2.211

McNamara, T. P., Rump, B., & Werner, S. (2003). Egocentric and geocentric frames of reference in memory of large-scale space. Psychonomic Bulletin & Review, 10(3), 589–595. 10.3758/BF03196519

Meilinger, T. (2008). The Network of Reference Frames Theory: A Synthesis of Graphs and Cognitive Maps. In C. Freksa, N. S. Newcombe, P. Gärdenfors, & S. Wölfl (Eds.), Spatial Cognition VI. Learning, Reasoning, and Talking about Space (Vol. 5248, pp. 344–360). Springer Berlin Heidelberg. 10.1007/978-3-540-87601-4_25

Meilinger, T., Riecke, B. E., & Bülthoff, H. H. (2014). Local and Global Reference Frames for Environmental Spaces. Quarterly Journal of Experimental Psychology, 67(3), 542–569. 10.1080/17470218.2013.821145

Miller, J., Watrous, A. J., Tsitsiklis, M., Lee, S. A., Sheth, S. A., Schevon, C. A., Smith, E. H., Sperling, M. R., Sharan, A., Asadi-Pooya, A. A., Worrell, G. A., Meisenhelter, S., Inman, C. S., Davis, K. A., Lega, B., Wanda, P. A., Das, S. R., Stein, J. M., Gorniak, R., & Jacobs, J. (2018). Lateralized hippocampal oscillations underlie distinct aspects of human spatial memory and navigation. Nature Communications, 9(1), 2423. 10.1038/s41467-018-04847-9

Moar, I., & Bower, G. H. (1983). Inconsistency in spatial knowledge. Memory & Cognition, 11(2), 107–113. 10.3758/BF03213464

Montello, D. R. (1993). Scale and multiple psychologies of space. In A. U. Frank & I. Campari (Eds.), Spatial Information Theory A Theoretical Basis for GIS (Vol. 716, pp. 312–321). Springer Berlin Heidelberg. 10.1007/3-540-57207-4_21

Moser, E. I., & Moser, M.-B. (2008). A metric for space. Hippocampus, 18(12), 1142–1156. 10.1002/hipo.20483

Mou, W., & McNamara, T. P. (2002). Intrinsic frames of reference in spatial memory. Journal of Experimental Psychology: Learning, Memory, and Cognition, 28(1), 162–170. 10.1037/0278-7393.28.1.162

Mou, W., McNamara, T. P., Rump, B., & Xiao, C. (2006). Roles of egocentric and allocentric spatial representations in locomotion and reorientation. Journal of Experimental Psychology: Learning, Memory, and Cognition, 32(6), 1274–1290. 10.1037/0278-7393.32.6.1274

Mou, W., Zhao, M., & McNamara, T. P. (2007). Layout geometry in the selection of intrinsic frames of reference from multiple viewpoints. Journal of Experimental Psychology: Learning, Memory, and Cognition, 33(1), 145–154. 10.1037/0278-7393.33.1.145

Muryy, A., & Glennerster, A. (2018). Pointing Errors in Non-metric Virtual Environments. In S. Creem-Regehr, J. Schöning, & A. Klippel (Eds.), Spatial Cognition XI (Vol. 11034, pp. 43–57). Springer International Publishing. 10.1007/978-3-319-96385-3_4

Muryy, A., & Glennerster, A. (2021). Route selection in non-Euclidean virtual environments. PLOS ONE, 16(4), e0247818. 10.1371/journal.pone.0247818

Nakaya, T. (1997). Statistical Inferences in Bidimensional Regression Models. Geographical Analysis, 29(2), 169–186. 10.1111/j.1538-4632.1997.tb00954.x

Nau, M., Navarro Schröder, T., Bellmund, J. L. S., & Doeller, C. F. (2018). Hexadirectional coding of visual space in human entorhinal cortex. Nature Neuroscience, 21(2), 188–190. 10.1038/s41593-017-0050-8

O’Keefe, J., & Dostrovsky, J. (1971). The hippocampus as a spatial map. Preliminary evidence from unit activity in the freely-moving rat. Brain Research, 34(1), 171–175. 10.1016/0006-8993(71)90358-1

O’Keefe, J., & Nadel, L. (1978). The hippocampus as a cognitive map. Clarendon Press; Oxford University Press.

Park, S. A., Miller, D. S., & Boorman, E. D. (2021). Inferences on a multidimensional social hierarchy use a grid-like code. Nature Neuroscience, 24(9), 1292–1301. 10.1038/s41593-021-00916-3

Peer, M., Ron, Y., Monsa, R., & Arzy, S. (2019). Processing of different spatial scales in the human brain. eLife, 8, e47492. 10.7554/eLife.47492

Richardson, A. E., Montello, D. R., & Hegarty, M. (1999). Spatial knowledge acquisition from maps and from navigation in real and virtual environments. Memory & Cognition, 27(4), 741–750. 10.3758/BF03211566

Rieser, J. J. (1989). Access to knowledge of spatial structure at novel points of observation. Journal of Experimental Psychology. Learning, Memory, and Cognition, 15(6), 1157–1165. 10.1037//0278-7393.15.6.1157

Rolls, E. T. (2023). Hippocampal spatial view cells, place cells, and concept cells: View representations. Hippocampus, 33(5), 667–687. 10.1002/hipo.23536

Ruginski, I. T., Creem-Regehr, S. H., Stefanucci, J. K., & Cashdan, E. (2019). GPS use negatively affects environmental learning through spatial transformation abilities. Journal of Environmental Psychology, 64, 12–20. 10.1016/j.jenvp.2019.05.001

Shelton, A. L., & McNamara, T. P. (1997). Multiple views of spatial memory. Psychonomic Bulletin & Review, 4(1), 102–106. 10.3758/BF03210780

Shelton, A. L., & McNamara, T. P. (2001). Systems of Spatial Reference in Human Memory. Cognitive Psychology, 43(4), 274–310. 10.1006/cogp.2001.0758

Shelton, A. L., & McNamara, T. P. (2004). Orientation and Perspective Dependence in Route and Survey Learning. Journal of Experimental Psychology: Learning, Memory, and Cognition, 30(1), 158–170. 10.1037/0278-7393.30.1.158

Stachenfeld, K. L., Botvinick, M. M., & Gershman, S. J. (2017). The hippocampus as a predictive map. Nature Neuroscience, 20(11), 1643–1653. 10.1038/nn.4650

Starrett, M. J., Stokes, J. D., Huffman, D. J., Ferrer, E., & Ekstrom, A. D. (2019). Learning-dependent evolution of spatial representations in large-scale virtual environments. Journal of Experimental Psychology: Learning, Memory, and Cognition, 45(3), 497–514. 10.1037/xlm0000597

Stella, F., Urdapilleta, E., Luo, Y., & Treves, A. (2020). Partial coherence and frustration in self-organizing spherical grids. Hippocampus, 30(4), 302–313. 10.1002/hipo.23144

Stevens, A., & Coupe, P. (1978). Distortions in judged spatial relations. Cognitive Psychology, 10(4), 422–437. 10.1016/0010-0285(78)90006-3

Taube, J. S. (2007). The Head Direction Signal: Origins and Sensory-Motor Integration. Annual Review of Neuroscience, 30(1), 181–207. 10.1146/annurev.neuro.29.051605.112854

Thorndyke, P. W. (1981). Distance estimation from cognitive maps. Cognitive Psychology, 13(4), 526–550. 10.1016/0010-0285(81)90019-0

Tobler, W. R. (1994). Bidimensional Regression. Geographical Analysis, 26(3), 187–212. 10.1111/j.1538-4632.1994.tb00320.x

Tversky, B. (1981). Distortions in memory for maps. Cognitive Psychology, 13(3), 407–433. 10.1016/0010-0285(81)90016-5

Tversky, B. (1992). Distortions in cognitive maps. Geoforum, 23(2), 131–138. 10.1016/0016-7185(92)90011-R

Tversky, B. (1993). Cognitive maps, cognitive collages, and spatial mental models. In A. U. Frank & I. Campari (Eds.), Spatial Information Theory A Theoretical Basis for GIS (Vol. 716, pp. 14–24). Springer Berlin Heidelberg. 10.1007/3-540-57207-4_2

Waller, D. A., & Nadel, L. (Eds.). (2013). Handbook of spatial cognition. American Psychological Association.

Waller, D., & Greenauer, N. (2007). The role of body-based sensory information in the acquisition of enduring spatial representations. Psychological Research, 71(3), 322–332. 10.1007/s00426-006-0087-x

Waller, D., & Hodgson, E. (2006). Transient and enduring spatial representations under disorientation and self-rotation. Journal of Experimental Psychology: Learning, Memory, and Cognition, 32(4), 867–882. 10.1037/0278-7393.32.4.867

Warren, W. H. (2019). Non-Euclidean navigation. The Journal of Experimental Biology, 222(Suppl 1), jeb187971. 10.1242/jeb.187971

Warren, W. H., Rothman, D. B., Schnapp, B. H., & Ericson, J. D. (2017). Wormholes in virtual space: From cognitive maps to cognitive graphs. Cognition, 166, 152–163. 10.1016/j.cognition.2017.05.020

Weisberg, S. M., Schinazi, V. R., Newcombe, N. S., Shipley, T. F., & Epstein, R. A. (2014). Variations in cognitive maps: Understanding individual differences in navigation. Journal of Experimental Psychology: Learning, Memory, and Cognition, 40(3), 669–682. 10.1037/a0035261

Widdowson, C., & Wang, R. F. (2022). Human navigation in curved spaces. Cognition, 218, 104923. 10.1016/j.cognition.2021.104923

Wolbers, T., & Hegarty, M. (2010). What determines our navigational abilities? Trends in Cognitive Sciences, 14(3), 138–146. 10.1016/j.tics.2010.01.001

Wolbers, T., & Wiener, J. M. (2014). Challenges for identifying the neural mechanisms that support spatial navigation: The impact of spatial scale. Frontiers in Human Neuroscience, 8. 10.3389/fnhum.2014.00571

